# Integrated multivariate analysis of transcriptomic data reveals immunological mechanisms in mice after *Leishmania infantum* infection

**DOI:** 10.1101/2020.10.09.333005

**Authors:** Génesis Palacios, Raquel Diaz-Solano, Basilio Valladares, Roberto Dorta-Guerra, Emma Carmelo

## Abstract

Transcriptional analysis of complex biological scenarios has been extensively used, even though sometimes results may prove imprecise or difficult-to interpret due to an overwhelming amount of information. In this study, a large-scale Real-time qPCR experiment was coupled to multivariate statistical analysis to describe the main immunological events underlying the early *L. infantum* infection in livers of BALB/c mice. High-throughput qPCR was used to evaluate the expression of 223 genes related to immunometabolism 1-, 3-, 5- and 10-days post infection. This integrative analysis showed strikingly different gene signatures at 1- and 10-days post infection, revealing progression of infection in the experimental model based on the upregulation of particular immunological response patterns and mediators. This approach addresses the challenge of integrating large collections of transcriptional data for the identification of candidate biomarkers in experimental models.

## INTRODUCTION

Visceral leishmaniasis (VL) is one of the clinical forms of leishmaniasis, caused mainly by the intracellular protozoan *Leishmania donovani* or *Leishmania infantum*. The murine animal model, mainly BALB/c and C57BL/6 strains, has been extensively used in the immunological study of this disease. When the infection is reproduced in the experimental model, immune response occurs in the main affected organs, liver and spleen. In the liver, the formation of cellular infiltrates or inflammatory granulomas occur around the infected macrophages. Kupffer cells (KC) are key in this process, since these are the main tissue macrophages that cover hepatic sinusoids and are the main target of infection by strains of *Leishmania* causing VL. Once the parasite is inoculated in the host, uptake of amastigotes by KC is rapid and completed within minutes (1). Nevertheless, uptake of *Leishmania* by macrophages depend on both host cell type and parasite species (2,3). For example, it was shown that the engulfment of intravenously inoculated promastigotes of *L. donovani* by neutrophils was faster than inoculated amastigotes (4).

When the infection occurs, the chemotaxis of NKT cells is promoted by the production of CXCL9: iNKT cells produce cytokines, including interferon-γ (IFN-γ), interleukin-4 (IL-4), and IL-17A, that stimulate innate immune responses such as neutrophil recruitment (5). CCL3, CCL2 and CXCL10 are initially secreted in the liver, possibly by infected KC (6); this generates the initial recruitment of monocytes and neutrophils, which are critical for effective control of parasite growth. At the first 2 hours post infection, studies in Jα281−/− mice indicated that only invariant NKT cells produced IFN-γ that was functionally relevant in this context (7). On the other hand, T lymphocytes are recruited towards the granuloma in formation, in response to various chemokine and cytokine stimuli after the recruitment of monocytes and neutrophils. Around the first week after infection, the recruitment of CD4+ and CD8+ T cells occurs with the upregulation of MHC class II by KC to facilitate interaction with CD4+ T cells (reviewed in (8)). The granuloma provides an inflammatory environment in which TNF and IFN-γ are secreted, promoting the production of reactive oxygen intermediates (ROI) and reactive nitrogen intermediates (RNI), by infected KCs for the destruction of the parasite (9). After the second week post infection, a peak of parasite load is reached in the cells that were infected. When 4 weeks have passed after the infection, mature granulomas are observed that provide an inflammatory environment for parasitic elimination, although complete healing is not achieved.

In the liver, acute infection occurs and eventually is resolved with little tissue damage (reviewed in (10)). Chronic infection is developed in the spleen, leading to splenomegaly, suppression of the immune response and persistence of the parasite (6).

Many studies have focused on the analysis of immunological response in the spleen in *Leishmania*-infected mice, (11–13), where many mediators play a role in M1/Th1 and M2/Th2 responses, but also markers of T reg response, Th17, and Tfh cells are involved (14). However, understanding the development of immunity in the liver, where infection develops shortly after inoculation, could lead to potential strategies to improve the elimination of the parasite in the spleen (15). The in-depth analyses of the processes that drive the early infection will provide a better insight on the immunological strategies that could control infection.

The methodology used in this study is based on High-Throughput-Real Time qPCR, for the amplification of 223 genes related to the immune response, lipid and carbohydrates metabolism. This is a highly sensitive and specific technology that allows a large-scale gene expression analysis and allows to correct experimental variations by normalization. Gene expression is also well-correlated with protein expression for numerous markers (such as IL12, IL10, IFN-γ, TLR2-4, TNF-α) in murine model (16). The transcriptional profile provides an approach of the mechanisms underlying the early *L. infantum* infection in murine model.

In this large-scale analysis, three different parameters were integrated: parasite burden in liver tissue, weight of organ and gene expression profile in the liver at different timepoints. This large dataset was streamlined using Principal Component Analysis (PCA) in order to identify a particular signature for some timepoints based on the contribution of each gene’s expression. PCA analysis revealed strikingly different gene-expression patterns between timepoints, unraveling the molecular processes taking place during early infection. In our experiments, at 1-day post inoculation, several genes were upregulated that are involved in different processes in response to IFN-γ, mediators at membrane level, but also in cell chemotaxis. However, at 10 dpi the gene signature was characterized as a more directed response with the upregulation of genes involved in *interleukin-12 signaling* pathway.

## RESULTS

### *Leishmania infantum* infection is established in liver 24 h post inoculation

As early as 24 h post inoculation, high parasitic burden was detected in liver tissue, indicating the infection is already established in this organ **(Figure 1A)**. This parasitic burden remains stable until 3 dpi; at 5 dpi, parasitic burden it jumps two orders of magnitude and remains elevated subsequently. Hepatomegaly in infected mice groups was not evident on days 1 and 3 post-infection, but small differences arise at 5 dpi and particularly at 10 dpi **(Figure 1B)**, which is in agreement with the leap in parasitic burden observed **(Figure 1A)**. Given our results, it seems clear that, although liver is already colonized 24 h after *L. infantum* inoculation, clinical signs such as patent inflammation appear a few days later in the infection in BALB/c mice.

**Figure 1.**
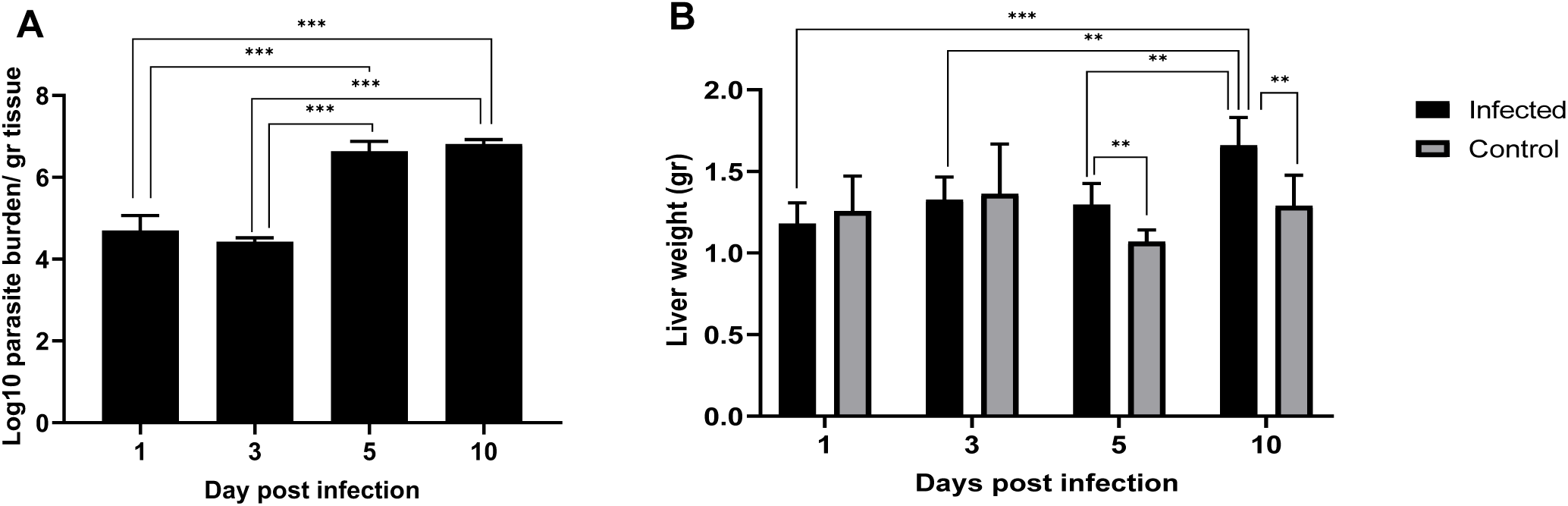
*Leishmania infantum* infection is established in liver 24 h post inoculation. (A) Evolution of parasite burden in liver. Mice were inoculated i.v. with 1 × 10^6^ promastigotes. Parasite load was determined 1 dpi, 3 dpi, 5 dpi and 10 dpi by limiting dilution assay and expressed as log^10^ of the average parasite load per gram of tissue. (B) Evolution of liver weight in infected (n = 24) vs. control mice (n = 24) over the course of infection. Error bars indicate SEM. One-way ANOVA was used to compare the differences in parasite burden between timepoints. Two-way ANOVA was used to compare the weight of liver between infected and control mice at each timepoint. Statistically significant differences are indicated (\**p* < 0.05; \*\**p* < 0.01, \*\*\**p* < 0.001).

### Global statistical approach of gene expression describes immunological events over time

Global analysis of transcriptome generated under different experimental conditions, has grown increasingly relevant in order to clarify key questions of intricate models like infection of target organs in leishmaniasis (11,13). With the aim of deciphering the most relevant aspects of the response against leishmaniasis developed in liver during the initial phases of infection with *L. infantum*, a massive real-time quantitative PCR analysis (more than 32000 qPCR reactions) was performed in BALB/c mice. The analysis mapped global changes in gene expression profiles of 223 genes of innate and adaptive immune response, prostaglandin synthesis, C-type lectin receptors, lipid metabolism and MAPK signaling pathway (**Supplementary file 1**). Candidate reference genes (n=9) and genes not expressed in any sample (n=14) were excluded for Principal Component Analysis (PCA). Global gene expression was evaluated from an immunometabolism perspective in infected vs. control livers, very early after infection: at 1-, 3-, 5- and 10-days after infection.

In order to identify gene expression profiles in this large dataset, PCA analysis was performed using Normalized Relative Quantity (NRQ) of all infected (n=24) and control (n=24) mice at all timepoints **(Figure 2 A)**. Four components were extracted with 60,84 % cumulative of total variance explained; Principal component 2 (PC2) and Principal component 3 (PC3) were used for subsequent analysis. PCA analysis showed that, 1-day post infection mice infected with *L. infantum* form a subset **(Figure 2 A**, indicated by a blue cloud). This was confirmed by two-way ANOVA (*p*< 0.001), having the score of PC2 as the response variable and time post infection (1-,3-,5- and 10-days post infection) and condition (infected or control), as factors. Therefore, as early as 24 hours post-inoculation, infected mice showed a clearly divergent gene-expression profile driven by the expression of genes correlated to PC2 as a response to infection by *L. infantum* **(Figure 2 B)**.

**Figure 2.**
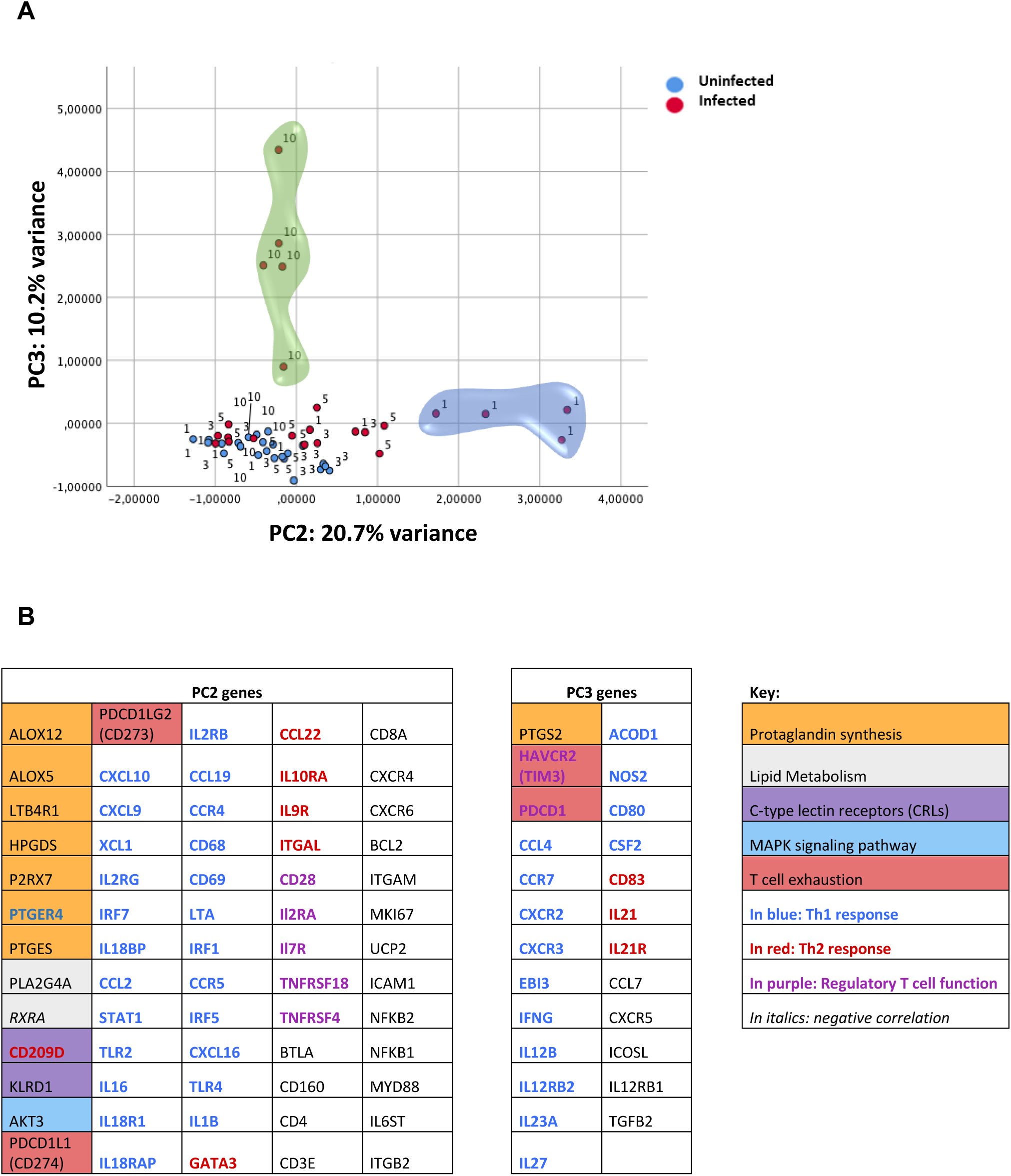
Global statistical approach of gene expression describes immunological events over time. qPCR was carried out on liver tissue from control and *L. infantum-*infected mice and at 1, 3, 5 and 10 dpi. (A) PCA plot. Principal components scores are shown on the X and Y axis, respectively, with the proportion of total variance related to that PC indicated. Each dot represents a mouse in the corresponding day and the color indicates the condition (red=infected or blue=control). The clouds highlight the cluster of 10 dpi-infected mice by PC3 (green cloud) and the cluster of 1 dpi-infected mice by PC2 (blue cloud). (B) List of PC2 and PC3 genes. “Key” indicates the classification of genes. The criteria to consider that a gene is correlated with a specific PC was a score ≥0.54 or ≤ - 0.54 of the factorial loadings. The following source data and figure supplements are available for figure 2: **Supplementary file 1**. List of TaqMan assays used for RT-qPCR analysis using QuantStudio™ 12K Flex Real-Time PCR System. **Figure 2- figure supplement 1**. Global statistical approach of gene expression - Unsupervised hierarchal clustering

PCA analysis also showed that *L. infantum* infected mice at 10 dpi form another subset **(Figure 2 A**, indicated by a green cloud). Using the score of PC3, we confirmed statistically significant differences between 10 dpi-infected and control mice (*p*<0.001), showing infected mice had also a clearly divergent gene-expression profile at this timepoint, associated to the expression of genes included in PC3 **(Figure 2 B)**.

Additionally, unsupervised hierarchal clustering of the NRQ values was applied and based on Euclidean distance and pairwise average-linkage was performed to visualize the relationships within this experimental dataset (**Figure 2- figure supplement 1**). Genes related to PC2 and PC3 were listed in **figure 2 B** and were classified depending whether they are involved in specific biological processes or biological pathways.

The list of genes correlated with PC2 include genes related to the immunological response triggered by infection, such as Th1/M1 markers, but also genes involved in Th2 response, prostaglandin synthesis, lipid metabolism, MAPK signaling pathway, regulatory T cell function, among others **(Figure 2 B)**. This indicates a very early activation (24 hours after infection) of diverse signals. In contrast, genes correlated with PC3 were not as numerous, and most of them are involved in Th1 response, suggesting the gene signature is tipping over to a more defined response at 10 dpi. The most striking findings will be described below.

In order to identify the global gene signatures that mark the difference between 1 dpi and 10 dpi-mice from the rest of mice, Gene Set Enrichment Analysis (GSEA) was applied to PC2 and PC3 correlated genes, to identify their most relevant signatures. With PC2 gene set, 1 dpi-mice were compared to the rest of mice, and with PC3 gene set, 10 dpi-mice were compared to the rest of mice.

With the set of genes correlated with PC2 (65 genes), 1058 gene sets were upregulated in 1 dpi-mice; only 59 gene sets were significant at FDR < 25% and 32 of them were significantly enriched at nominal *p*-value < 1%.

With the set of genes correlated with PC3 (24 genes), 681 gene sets were upregulated in 10 dpi-mice; 71 gene sets were significant at FDR < 25% and 28 gene sets were significantly enriched at nominal *p-*value < 1%.

In order to identify which functional categories are significantly over-represented by PC2 or PC3 genes, whether they are overlapping or not and how strong is the connection between them in terms of similarity coefficient, an enrichment map was constructed using Cytoscape 3.8.0, including both analysis with PC2 genes (dataset 1) and PC3 genes (dataset 2) with a total of 19 gene sets filtered with the parameters FDR < 0.25 and *p-*value < 0.001 **(Figure 3. A B**).

**Figure 3.**
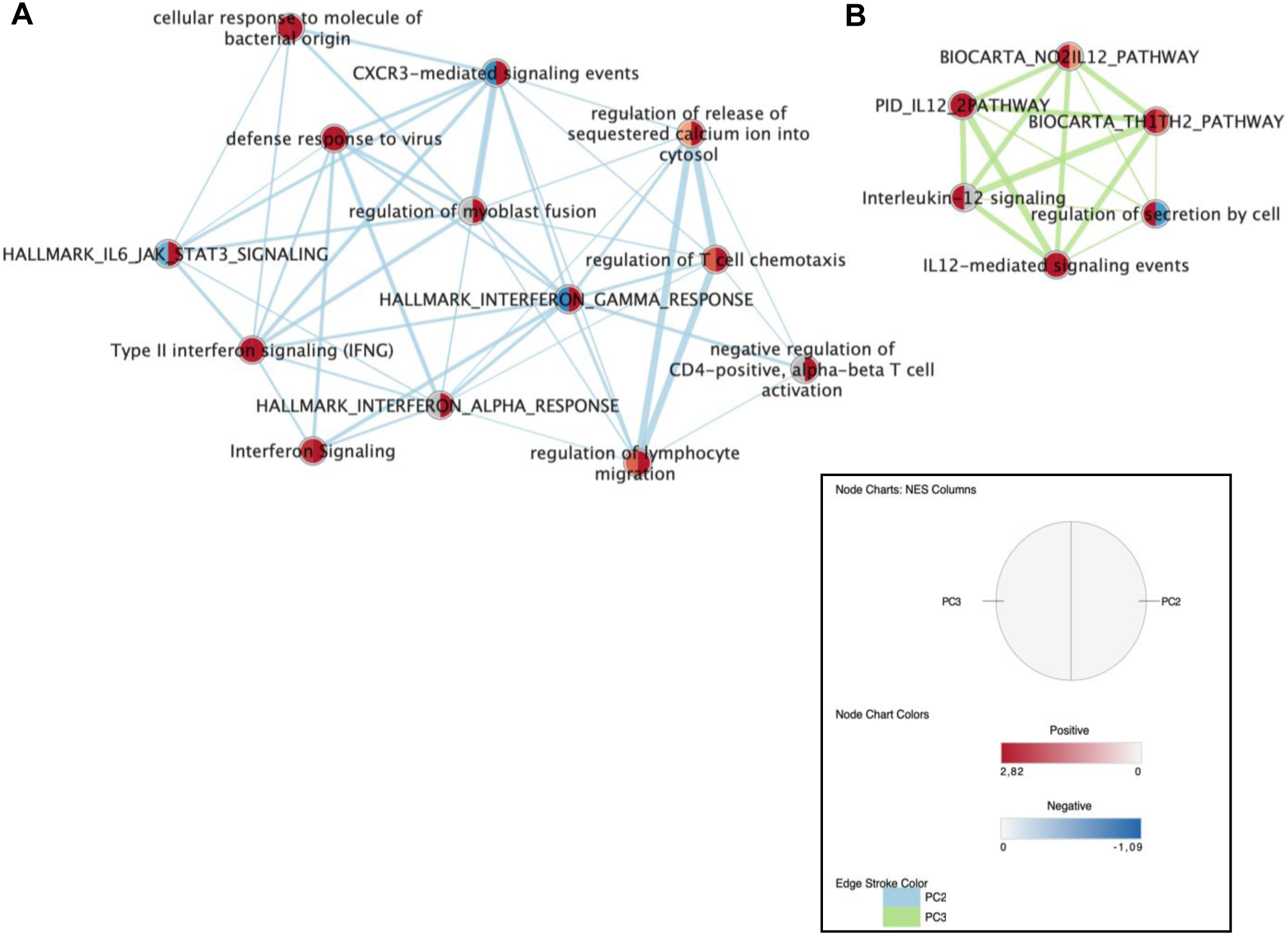
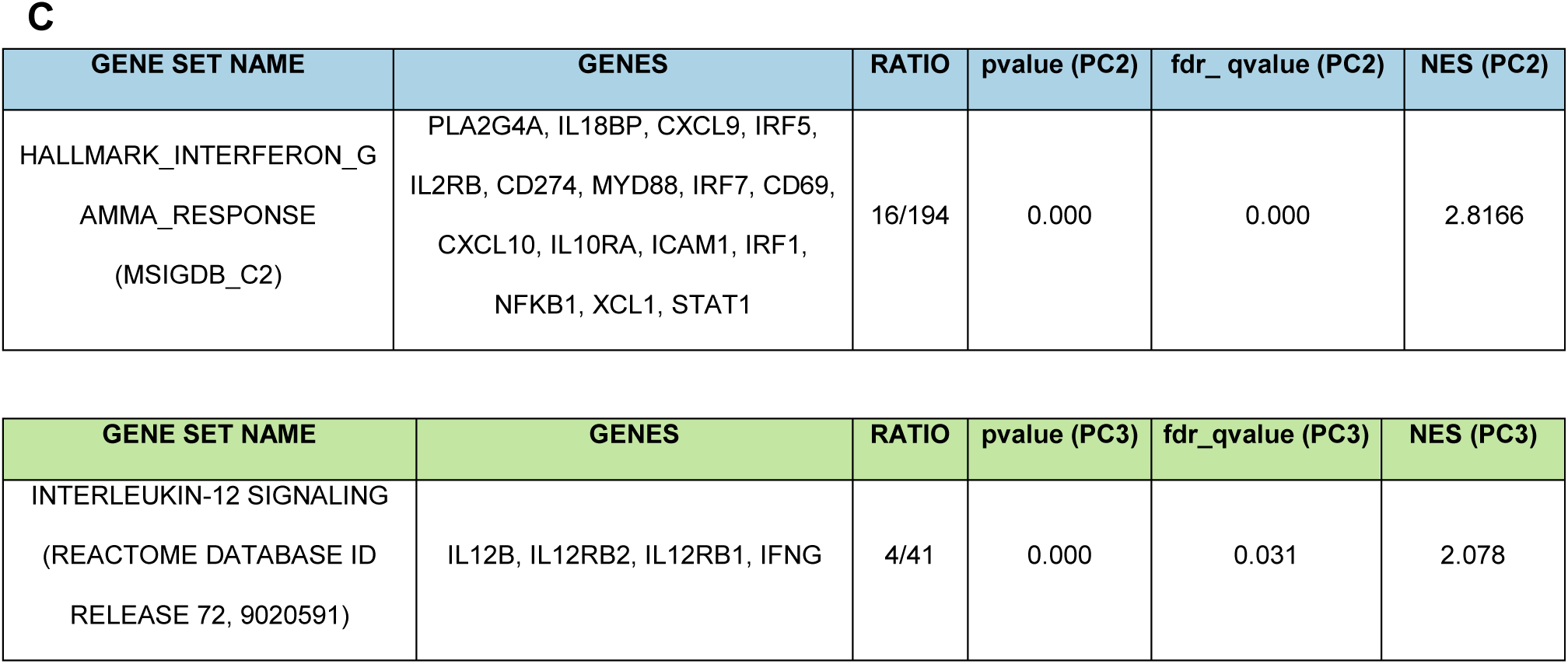
Enrichment Map combining both analysis with PC2 genes (dataset 1) and PC3 genes (dataset 2). **(A)** Group of nodes with blue line connections represents gene sets enriched by PC2 genes; **(B)** group of nodes with green line connections represents gene sets enriched by PC3 genes. Parameters to filter these nodes/gene sets applied in Cytoscape: *p*-value < 0.001 and FDR < 0.25. Node color indicates NES value (Normalized Enrichment Score), half right side: NES of PC2 enrichment analysis (deeper red color indicates upregulation in 1 dpi-mice, deeper blue color indicates downregulation in 1 dpi-mice); half left side: NES value of PC3 enrichment analysis (deeper red color indicates upregulation in 10 dpi-mice, deeper blue color indicates downregulation in 10 dpi-mice). Connection lines thickness represents similarity coefficient between nodes. **(C)** Central functional categories significantly upregulated by PC2 or PC3 genes in enrichment map (NES value > 0 and FDR < 0.05).

PC2 enrichment analysis resulted in many more gene sets enriched (*p*-value < 0.001 and FDR < 0.25) than PC3 enrichment analysis **(Figure 3. A B)**. Remarkably, gene sets enriched in 10 dpi-mice show strong association (signaled by thick green lines), indicating that all those categories are more connected among them than with gene sets enriched in 1 dpi-mice (blue lines). Considering functional categories which are significantly upregulated by PC2 or PC3 genes, we could identify the inherent processes in 1 dpi-mice or 10 dpi-mice that constitute the differences of this biological group in contrast with the others. This is the case for HALLMARK_INTERFERON_GAMMA_RESPONSE: it is strongly upregulated by PC2 and inhibited by PC3 genes; similarly, INTERLEUKIN-12 SIGNALING functional category is specifically upregulated by PC3 and was not enriched by PC2 genes (**Figure 3 C**).

The relevance and specificity of these strongly enriched functional categories is shown in **Figure 4**. Most genes of “HALLMARK_INTERFERON_GAMMA_RESPONSE”, were upregulated in 1 dpi-mice group but not in the rest of the mice (**Figure 4. A**). Similarly, “INTERLEUKIN-12 SIGNALING” genes were significantly upregulated only in 10 dpi-mice (**Figure 4. B**), revealing that although at 1 dpi many genes are upregulated in response to IFNG, the remarkable upregulation of IFNG itself is demonstrated at 10 dpi, with the upregulation of *Ifng, Il12b, Il12rb1* and *Il12rb2*.

**Figure 4.**
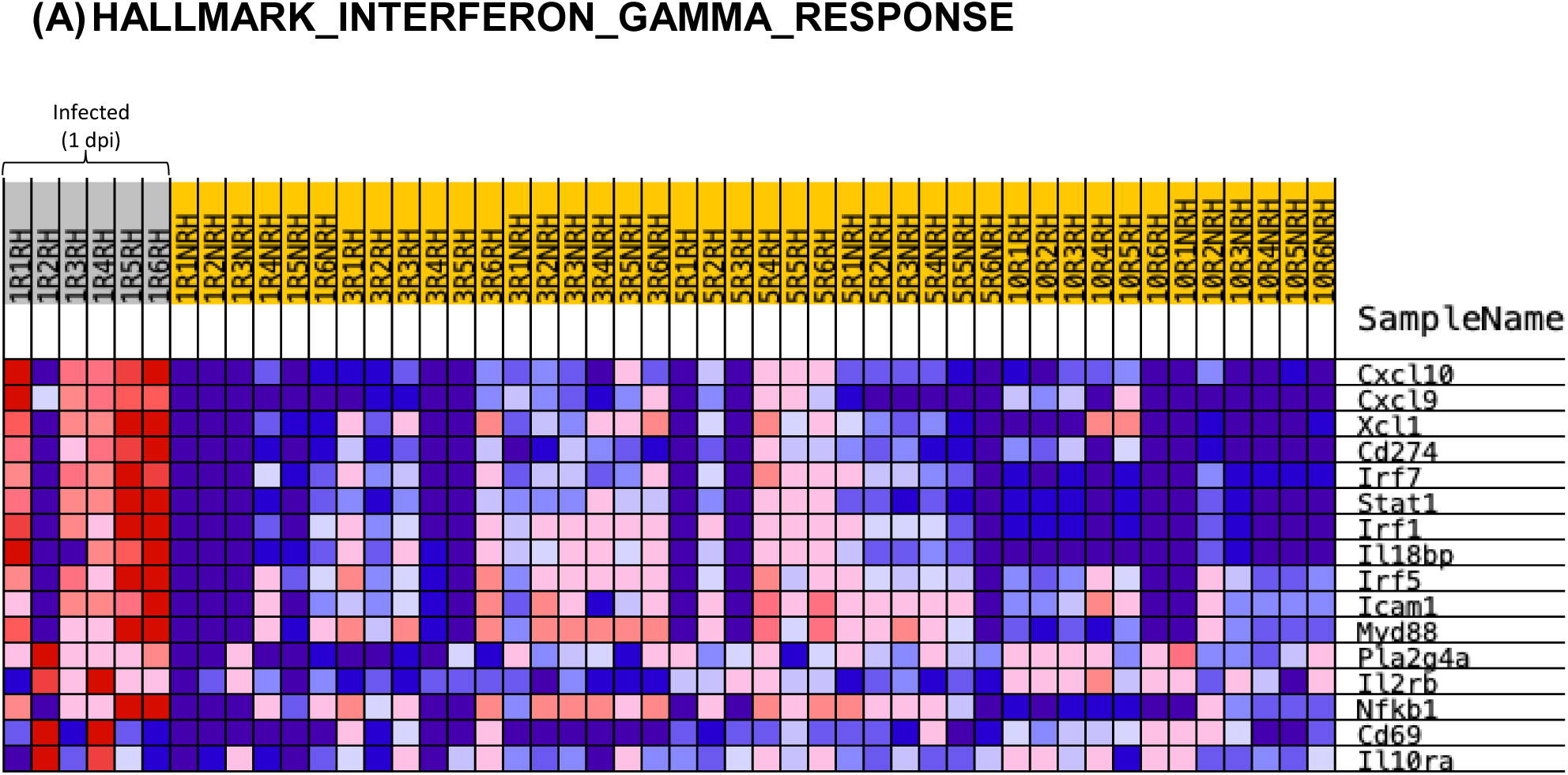
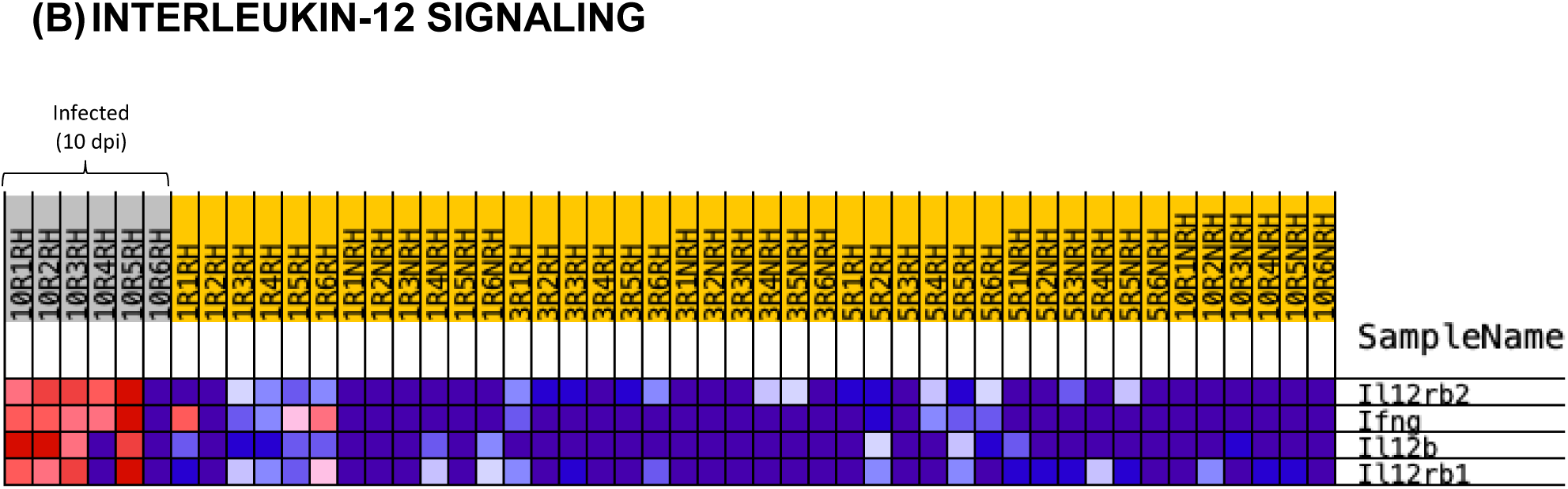
Gene set enrichment analysis (GSEA) identified the main enriched gene sets in 1 dpi-mice and 10-dpi mice. Genes are ordered by tTest ratio according to their differential expression. Normalized Relative Quantities (NRQ) are represented as colors, where the range of colors (red, pink, light blue, dark blue) shows the range of expression values (high, moderate, low, lowest). **(A) HALLMARK_INTERFERON_GAMMA_RESPONSE heatmap**. Genes upregulated in response to IFNG. This gene set was upregulated in 1 dpi-mice (PC2 dataset). Upregulation of these genes in 1 dpi -infected mice (left side) and the rest of mice is shown. Each mouse is identified with a respective code. **(B) INTERLEUKIN-12 SIGNALING heatmap**. This gene set was upregulated in 10 dpi-mice (PC3 dataset). Upregulation of these genes in 10 dpi -infected mice (left side) and the rest of mice is shown. Each mouse is identified with a respective code.

According to all these analyses, very early after parasite inoculation, diverse pathways and processes involved in immunological activation are displayed (1 dpi, PC2), although most of them are related to interferon signaling, suggesting a general non-specific activation of immune responses **(Figure 3. A)**. In contrast, at 10 dpi the main functional categories upregulated are related to interleukin-12 signaling (**Figure 3. B)**. These will be described in a more detailed way in independent Functional enrichment analysis using either PC2 or PC3 genes.

### Early inflammatory responses are activated in livers of *L. infantum* infected mice 1 day post inoculation

Enrichment analysis was performed using PC2 correlated genes, and an enrichment map **(Figure 5. A)** was generated to visualize the main functional categories, included in **Figure 5. B**. Thirteen of the gene sets showed *p*-value < 0.001 and FDR < 0.25, and five of these functional categories had the highest NES value (Normalized Enrichment Score, used to compare analysis results across gene sets (17)) and lowest FDR, therefore these categories were used to perform leading edge analysis (**Figure 5 C**). This analyses identifies the core that accounts for the gene set’s enrichment signal (17). It is notable the overrepresentation of some annotations such as HALLMARK_INTERFERON_GAMMA_RESPONSE, that showed the major connections among other annotations and the lowest FDR. This annotation connects to some others related to interferon signaling and to cell chemotaxis. (**Figure 5 A**).

**Figure 5.**
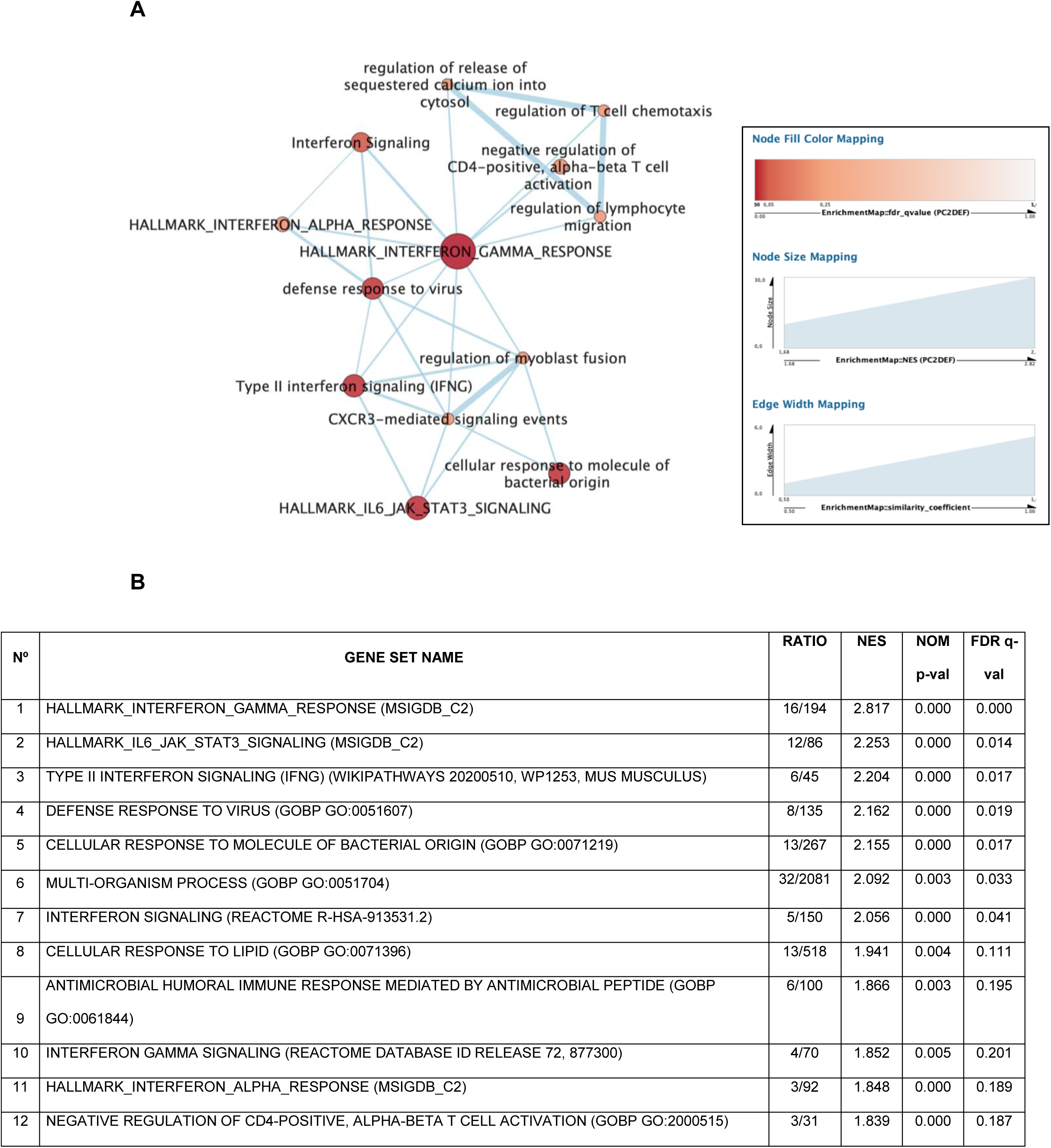
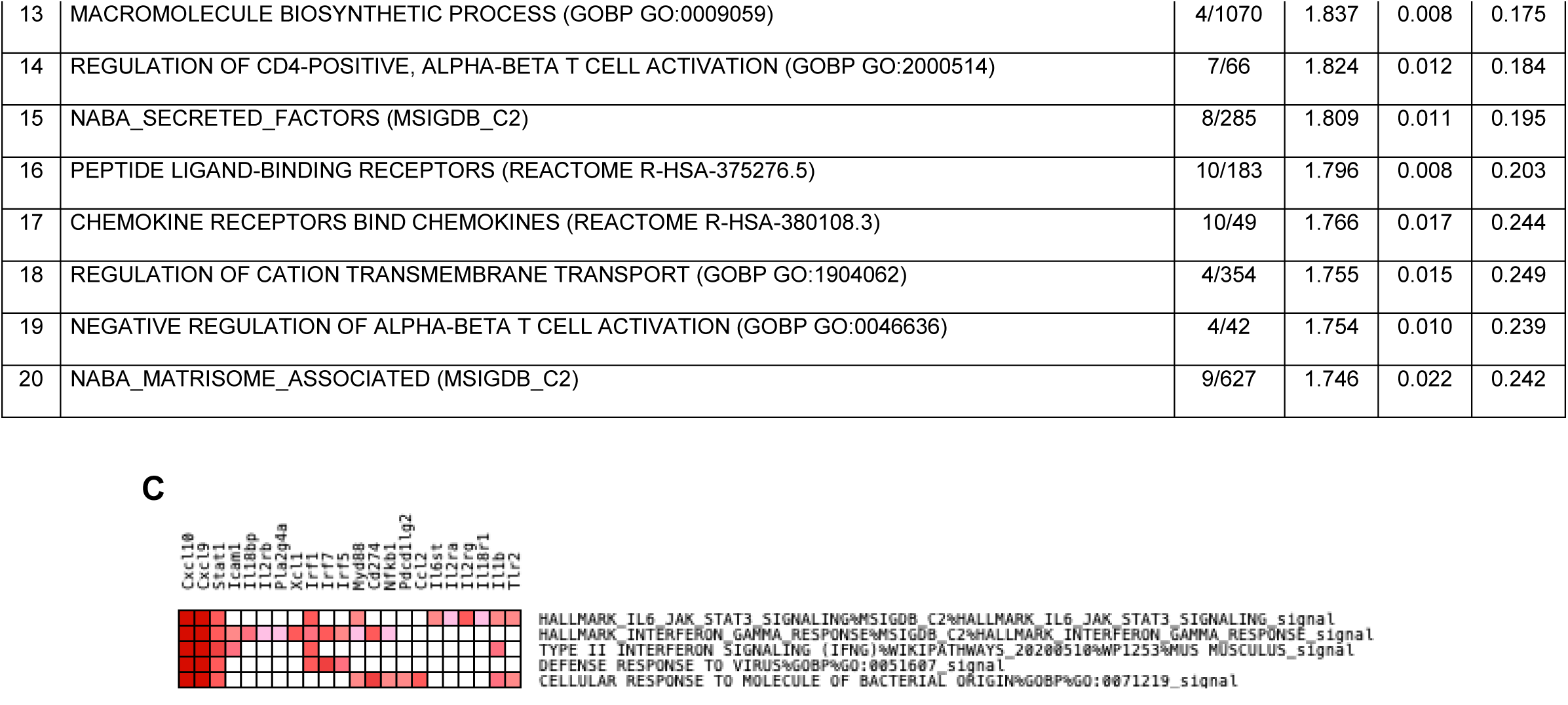
(A) Enrichment Map with PC2 correlated genes. Parameters to filter these nodes/gene sets applied in Cytoscape: *p*-value < 0.001 and FDR < 0.25. Intensity of node color fill represents FDR value (more intense red color indicates lower FDR); node size corresponds to the NES value. Width of connection lines represents similarity coefficient between nodes. (**B**) **Top 20 gene sets enriched in 1 dpi-mice**. Results of the enrichment analysis performed in GSEA software, the top 20 gene sets enriched with the highest NES values and with the lowest FDR. **(C) Leading edge analysis heatmap-PC2 dataset**. The analysis was performed in GSEA with the Top 5 gene sets enriched (with the highest NES value and lowest FDR value). The heat map shows the (clustered) genes in the leading edge subsets. In the heatmap, expression values are represented as colors, where the range of color (red, pink) shows the range of expression values (high, moderate).

Results of leading edge analysis remark the importance of *Cxcl10* and *Cxcl9* (**Figure 5 C**). The upregulation of these two markers indicates that at 1 dpi, neutrophils, one of the first group of leukocytes to arrive at the site of infection (18), would be generating chemotactic signals to NK cells.

With the aim of analyzing in detail the relations among PC2 genes, a functional enrichment analysis was performed in Cytoscape (**Figure 6**): Markov cluster algorithm was performed to determine clusters, with a topological clustering algorithm to partition the network based on the interactions, in that way we can explore the functional interactions of the network.

**Figure 6.**
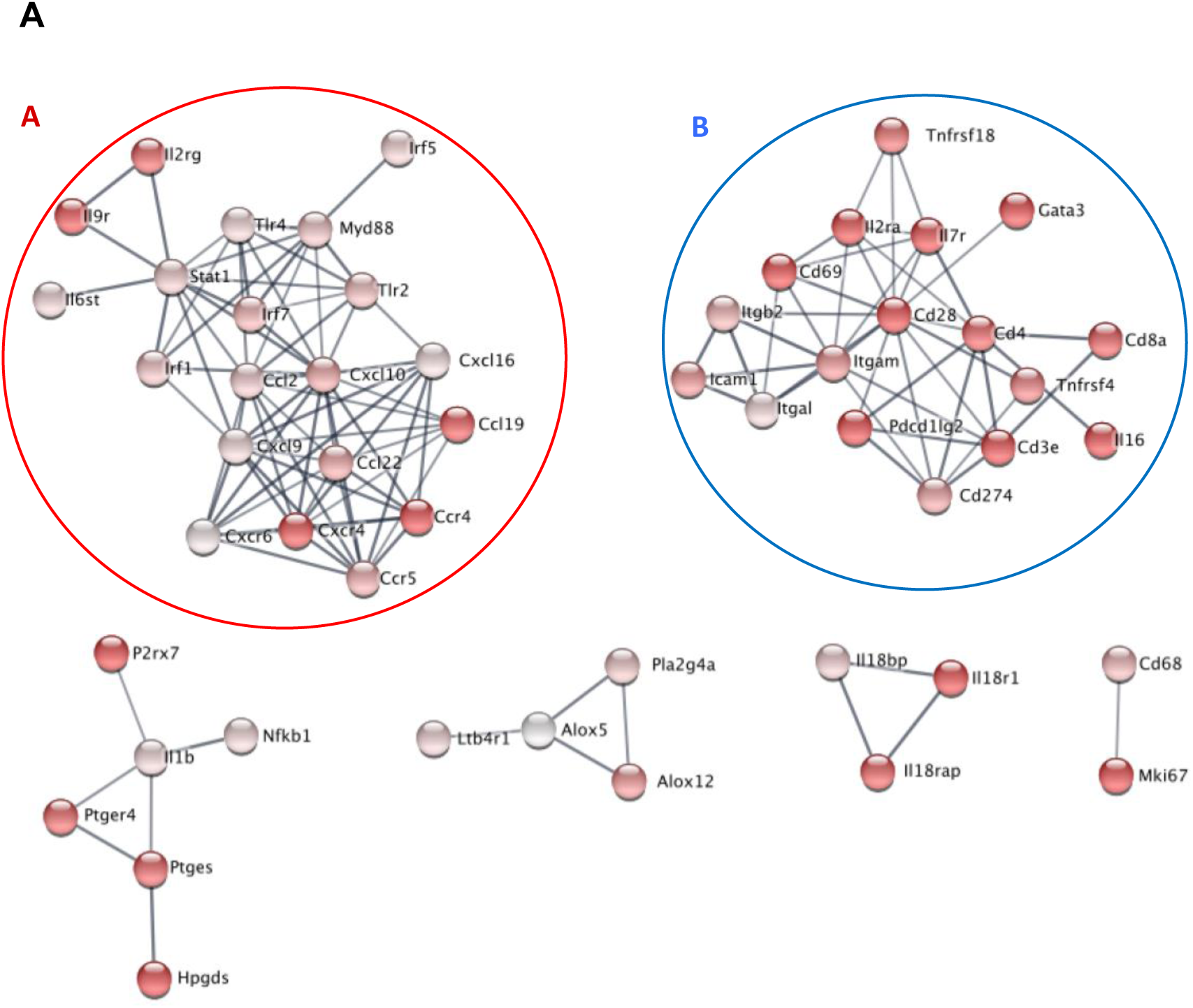
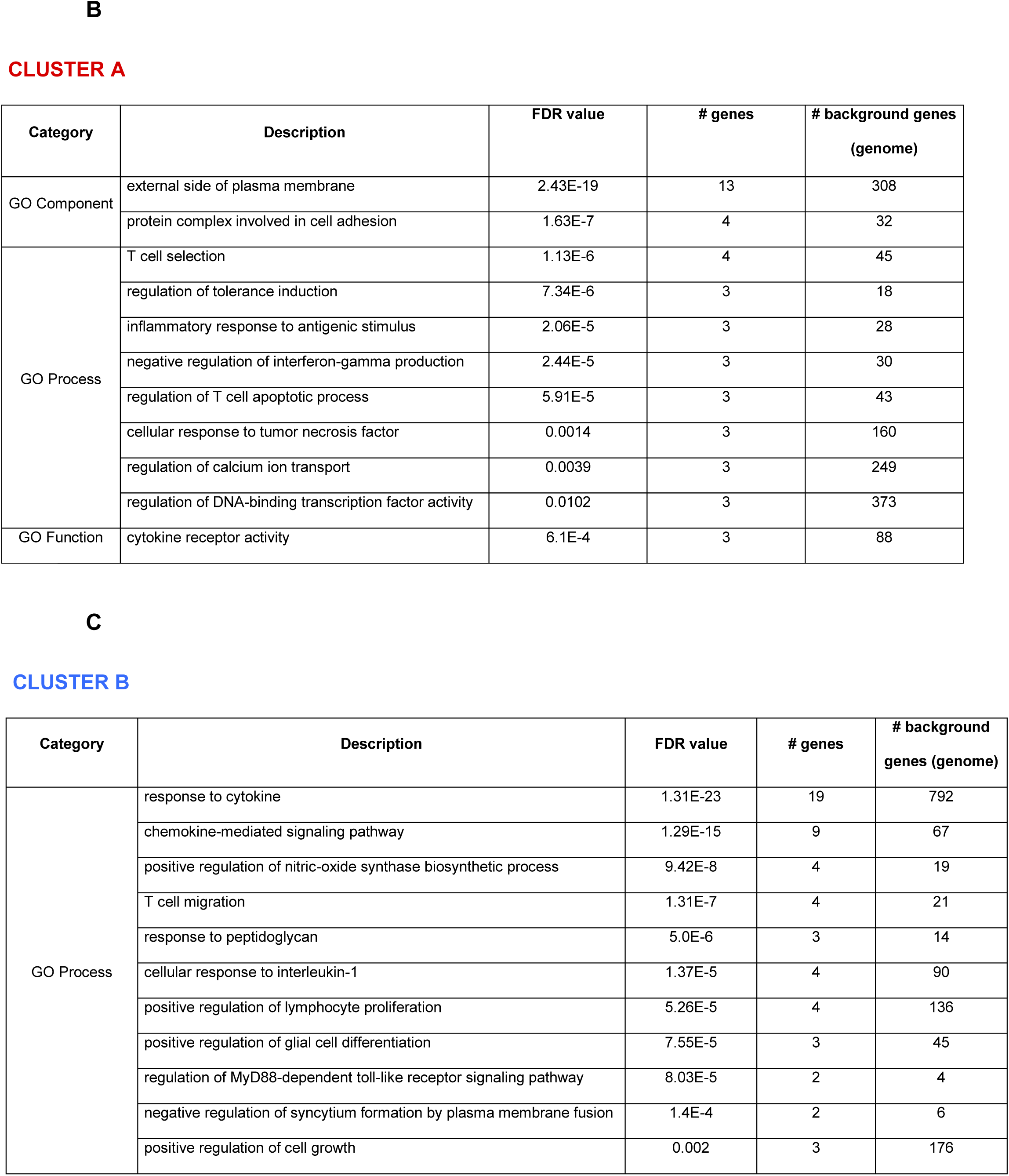
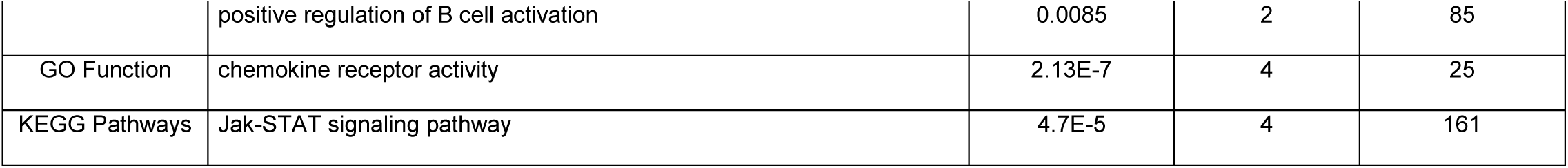
Functional enrichment analysis performed using STRING database. **(A)** STRING interaction network. Minimum required interaction score is medium confidence (0.7). Intensity of node color indicates correlation with PC2. Parameter of Markov Cluster Algorithm: Granularity parameter (inflation value): 5. **(B)** Cluster A. STRING functional enrichment. Similar terms were fused to reduce redundancy (Redundancy cut off= 0.25). **(C)** Cluster B STRING functional enrichment. Similar terms were fused to reduce redundancy (Redundancy cut off= 0.25). The following figure supplements are available for figure 6: **Figure 6- figure supplement 1**. Genes including in GO Component: External side of the plasma membrane.

In **Figure 6 A**, the STRING database was used to construct a protein–protein interaction (PPI) network. Within the genes correlated with PC2, STRING analysis showed two clusters, A and B, indicating the genes whose expression is strongly associated **(Figure 6. Cluster A and B)** and correlated with PC2 (indicated by intensity of node color).

Except RXRA, all genes were positively correlated with PC2. In other words, at 1-day post infection (and only at this time of infection), the particular gene-expression signature observed in infected mice is dependent on upregulation of all PC2 genes but down-regulation of RXRA (**Figure 10- figure supplement 1**).

In Cluster A, the annotation with the lowest FDR was *“External side of the plasma membrane*” (FDR value=2.43E-19, **Figure 6 B)** and includes genes that are playing a key role in livers of infected mice at 1dpi: *Il7r, Cd274, Cd4, Cd28, Il2ra, Tnfrsf4, Cd69, Cd8a, Itgam, Icam1, Tnfrsf18, Cd3e, Itgal*. The upregulation of these genes at 1 dpi (**Figure 6- figure supplement 1**), most of them coding for receptors and cell adhesion molecules, suggests the activation of early innate responses upon *L. infantum* infection.

In the main cluster of genes correlated with PC2 that lead towards the differentiation of 1 dpi-infected mice, *Cd69, an* iNKT cell marker, strikes as strongly upregulated. Outside this cluster we found other NK cell markers such as *klrd1*, which was also upregulated at this timepoint (**Figure 7**).

**Figure 7.**
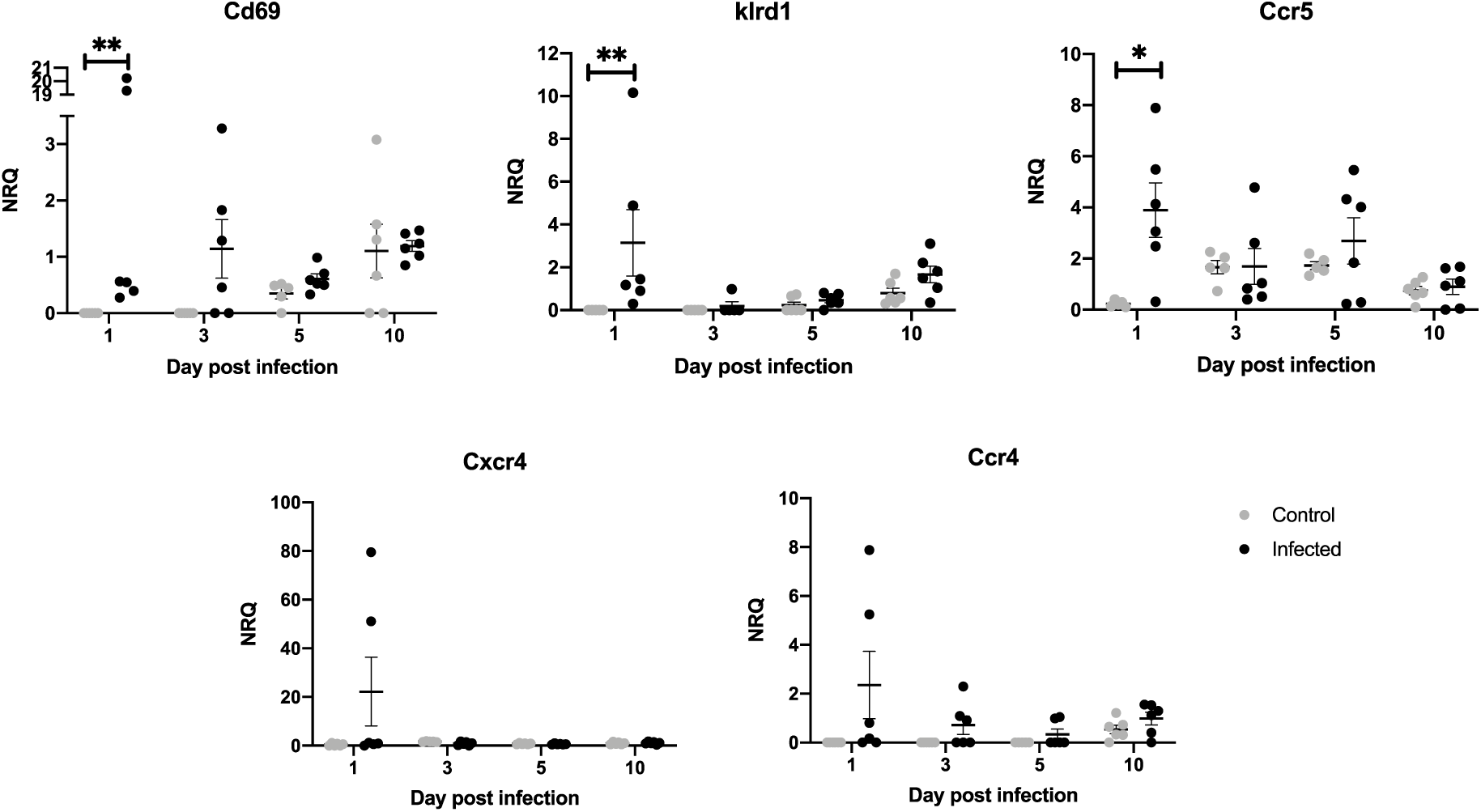
mRNA expression of genes expressed in NK or iNKT cells at 1 day post infection. Data are shown as the mean and SEM of the Normalized Relative Quantity values (NRQ). NRQ of infected mice (black symbols) and control mice (gray symbols) are represented in scatter plot. Differences in NRQ values between non-infected (control) and infected animals were analyzed by two tail Mann-Whitney test or two tail unpaired t-test *p<0.05; **p<0.01.

Among the GO Biological processes overrepresented by genes in cluster A, “*T cell selection*” stands out, with the participation of *Cd4, Cd28, Cd3e* and *Gata3*. The upregulation of those genes is also related to T cell activation, via CD28-CD86 costimulatory signaling. The central role of *Cd28* (**Figure 6 A)** in cluster A, and the upregulation of *Cd86* at 1 dpi (**Figure 11- figure supplement 1**), indicates T cell activation could be promoted at 1 dpi.

Interestingly, GO processes “regulation of tolerance induction” (*Cd274, Il2ra, Cd3e*) and “negative regulation of interferon-gamma production” (*Cd274, Gata3, Cd273*) are overrepresented by genes in cluster A (**Figure 6 C**), and both PDCD1 ligands were upregulated at 1 dpi: *Pd1l1* (or *Cd274*) and *Pd1l2* (or *Cd273*) (**Figure 8**). Despite the overrepresentation of those inhibitory categories in our study, *Pdcd1* did not show differential expression at this timepoint between infected vs control mice (**Figure 8**) and there are evidences of upregulation of many Th1 markers at 1 dpi.

**Figure 8.**
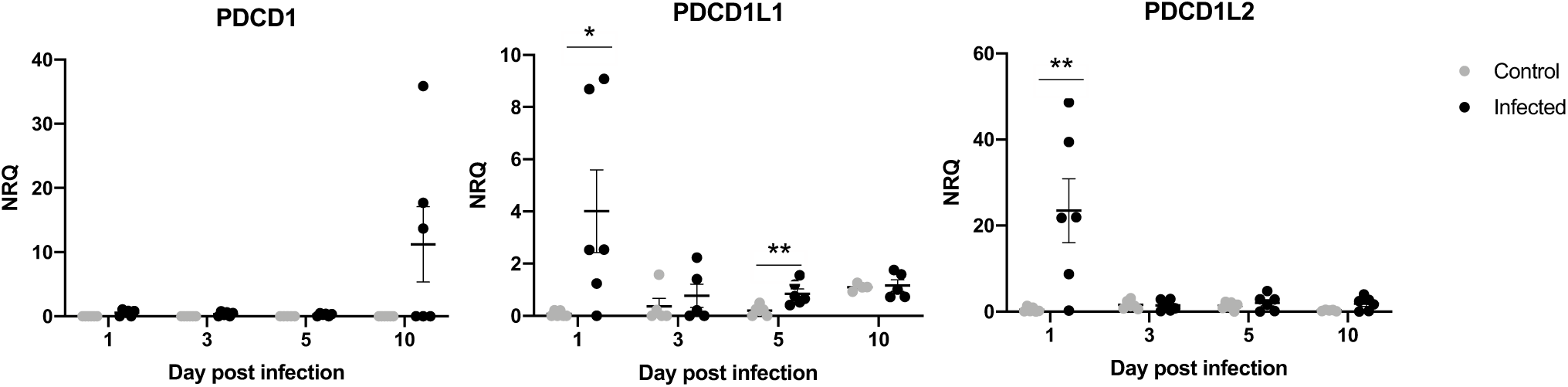
mRNA. expression of some T cells exhaustion genes. Data are shown as the mean and SEM of the Normalized Relative Quantity values (NRQ). NRQ of Infected mice (black symbols) and control mice (gray symbols) are represented in scatter plot. Differences in NRQ values between non-infected (control) and infected animals were analyzed by two tail Mann-Whitney test or two tail unpaired t-test *p<0.05; **p<0.01.

Cluster B shows another collection of genes well associated with each other **(Figure 6 A, Cluster B)**, but not as strongly correlated with PC2 (indicated by pale node color); the over-representation in GO Biological processes and KEGG pathways in cluster B is shown in **Figure 6 C**. One of the most remarkable annotations in cluster B is “chemokine-mediated signaling pathway” (*Ccl2, Ccl22, Cxcl10, Cxcr4, Cxcr6, Ccr4, Ccl19, Ccr5, Cxcl9*), that is one of the earliest outcomes of *Leishmania* infection described both *in vitro* and *in vivo* (6,19–22). The upregulation of *Tlr2* and *Tlr4* at 1 dpi (**Figure 6 A**) suggests that the chemokine response could be triggered by upstream signalling by Toll-like receptors.

It is also notable the over-representation of prostaglandin biosynthesis by HPGDS (PTGDS2) and PTGES **(Figure 6 A)**. These genes are included in *Arachidonic acid metabolism pathway* (mmu0059, FDR=0.002).

The *eicosanoid biosynthetic process* was over-represented by *Alox12, Alox5, Pla2g4a* (GO.0046456, FDR= 4.36E-6), and this group of genes was also well connected (**Figure 6**). These, and the genes encoding for *inflammatory response* (GO.0006954, FDR= 3,29E-27): *Ccl2, Itgb2, Lta, Alox5, Il18rap, Xcl1, Il2ra, Il1b, Tlr2, Tnfrsf4, Ccl22, Myd88, Tlr4, Cxcl10, Ltb4r1, Cxcr6, Ccr4, Itgam, Icam1, P2rx7, Tnfrsf18, Ptges, Ccl19, Gata3, Il18r1, Ccr5, Nfkb2, Cxcl9*, suggest that the first signals for inflammation of the liver have been issued, although hepatomegaly is not yet evident.

### *L. infantum* infection induces Th1/M1 responses through interleukin-12 signaling and positive regulation of interferon-gamma production at 10 days post infection

PCA allowed the discrimination of infected mice 10 days post inoculation **(Figure 2 A)**. This differentiation owes to the upregulation of some genes at that timepoint **(Figure 9)**, that are the genes correlated with PC3 in the Principal Component Analysis.

**Figure 9.**
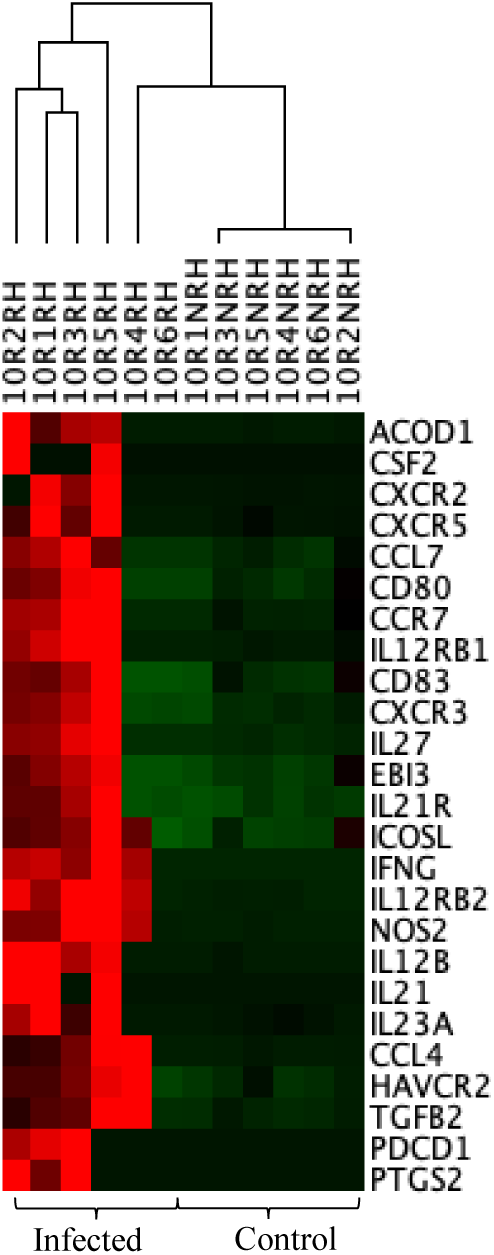
Unsupervised hierarchal clustering of 10 dpi- infected and control mice including genes correlated with PC3 was applied and based on Euclidean distance and pairwise average-linkage. Differential expression of selected genes in 10 dpi -infected mice (left side) and control mice (right side) is shown. Each mouse is identified with a respective code. Genes down regulated during infection are shown in green and upregulated genes are in red.

Enrichment map was constructed using PC3 correlated genes (**Figure 10 A)** to visualize the main functional categories, using the same stringent parameters stablished for the enrichment map with PC2 genes (**Figure 5 A**). Under these conditions, only 5 gene sets were upregulated by PC3 genes (highest NES values and lowest FDR), 4 of them related to IL-12 signaling pathway (PID_IL12_2PATHWAY, BIOCARTA_NO2IL12_PATHWAY, IL12-MEDIATED SIGNALING EVENTS, INTERLEUKIN-12 SIGNALING). The leading-edge analysis showed the relevance of *Il12b, Il12rb1, Il12rb2* and *Ifng*, all of them shared by these 4 pathways (**Figure 10 C**). At 10 days post infection, the upregulation of markers in **Figure 10 C**, suggests that the upregulation of *Ifng* could be a consequence of the activation of the IL12 signaling.

**Figure 10.**
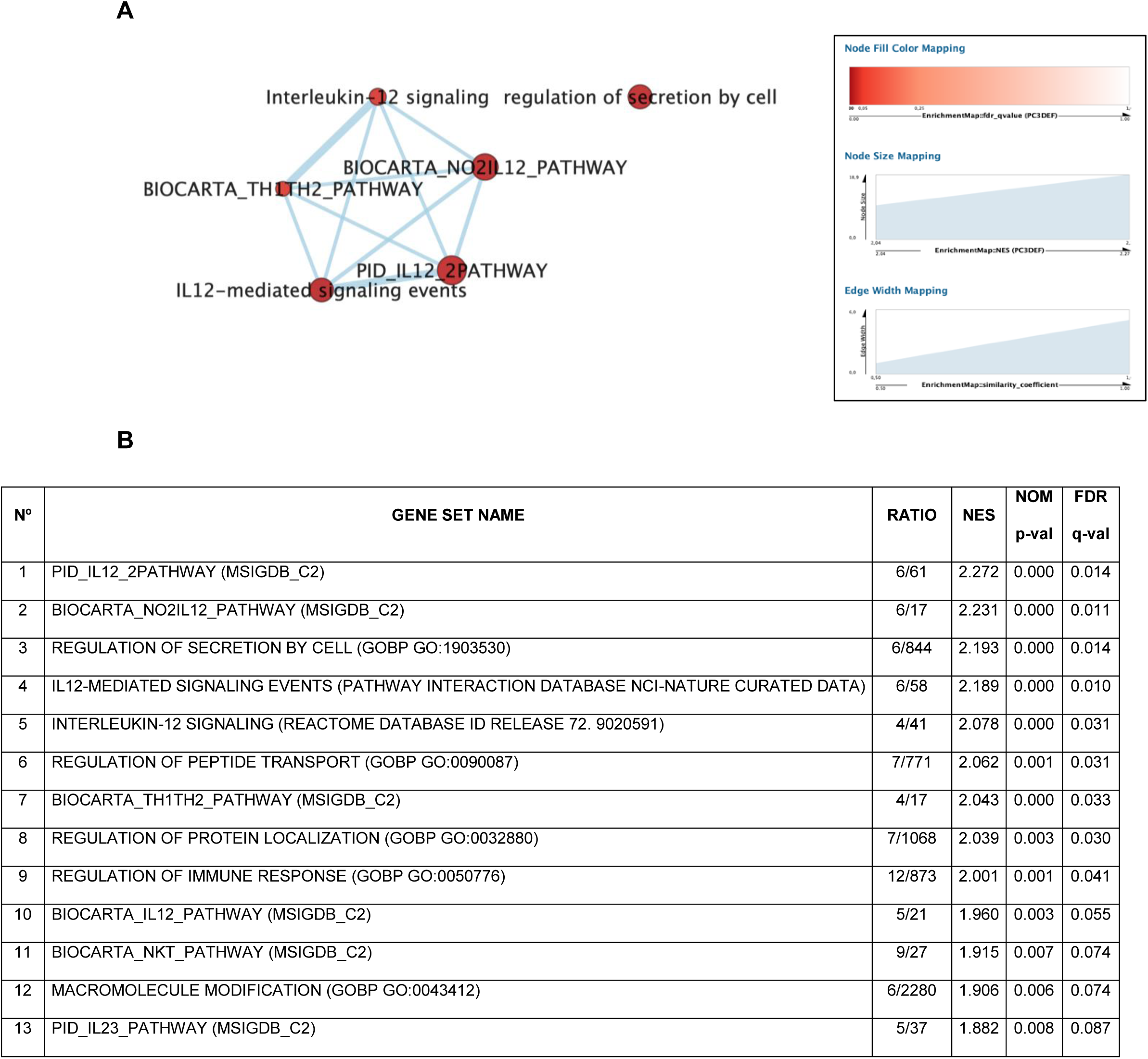
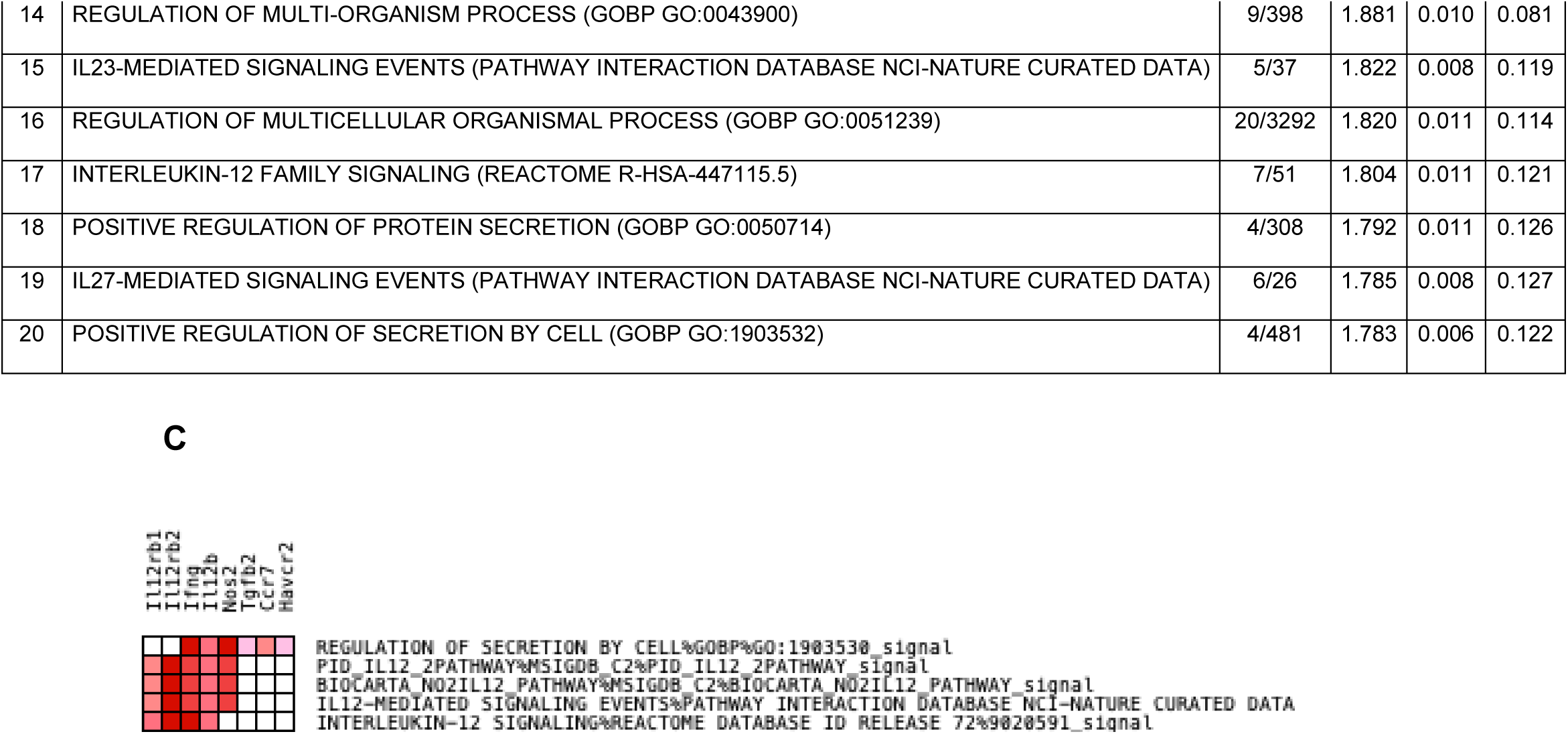
(A) Enrichment Map with PC3 correlated genes shows the relevance of IL-12 signaling pathway at 10 dpi. Parameters to filter these nodes/gene sets applied in Cytoscape: *p*-value < 0.001 and FDR < 0.25. Intensity of node color fill represents FDR value (more intense red color indicates lower FDR); node size corresponds to the NES value. Width of connection lines represents similarity coefficient between nodes. **(B) Top 20 gene sets enriched in 10 dpi-mice**. Results of the enrichment analysis performed in GSEA software, the top 20 gene sets enriched with the highest NES values and with the lowest FDR. **(C) Leading edge analysis heatmap-PC3 dataset**. The analysis was performed in GSEA with the Top 5 gene sets enriched (with the highest NES value and lowest FDR value). The heat map shows the (clustered) genes in the leading edge subsets. In the heat map, expression values are represented as colors, where the range of color (red, pink) shows the range of expression values (high, moderate).

To analyze the relationship among genes correlated with PC3, functional enrichment analysis was performed using STRING database (**Figure 11**) (24). From the STRING Interaction Network, Markov clustering was conducted, and one cluster is observed with most genes strongly correlated with PC3. All these genes were upregulated at 10 dpi (**Figure 9**).

**Figure 11.**
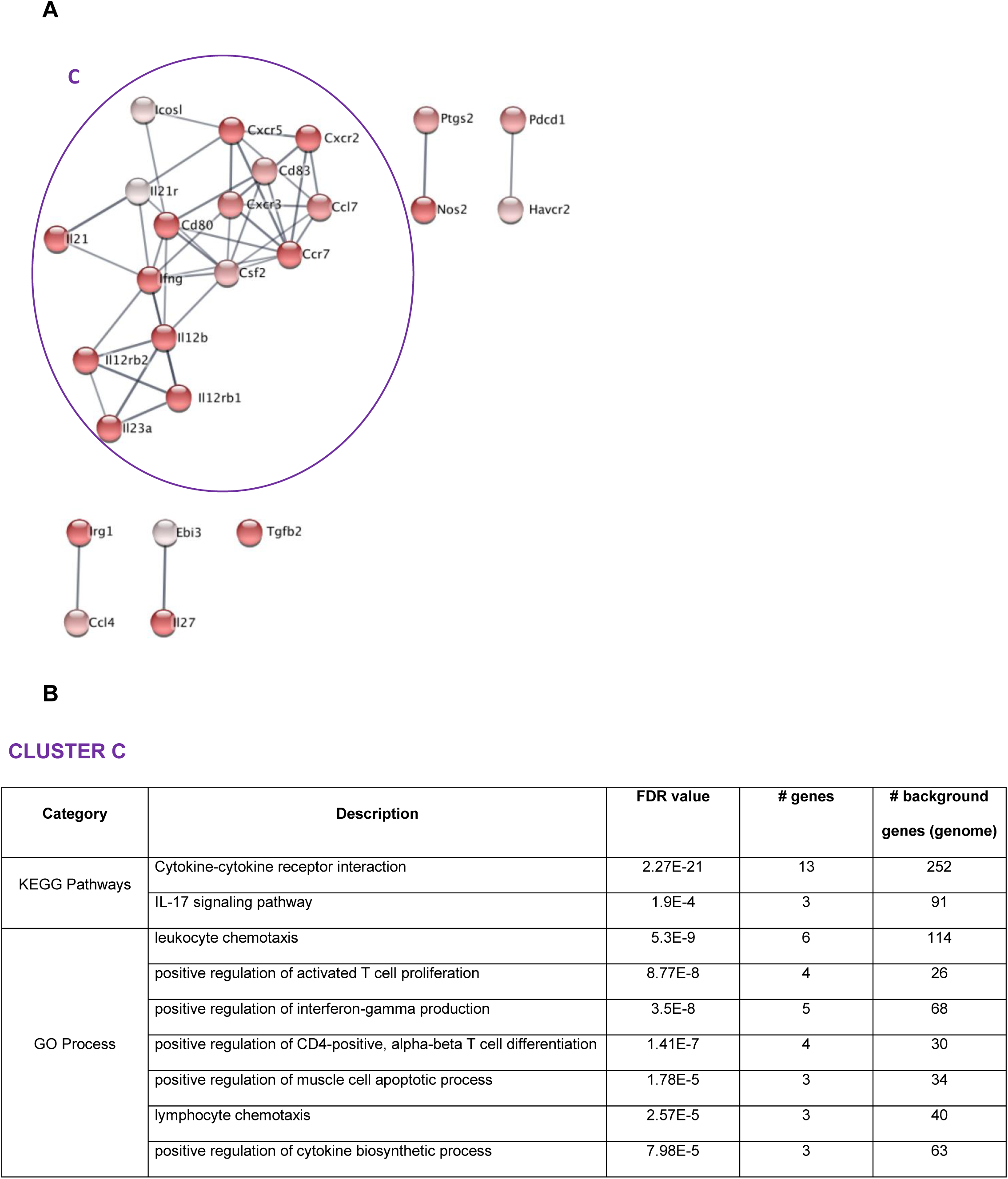
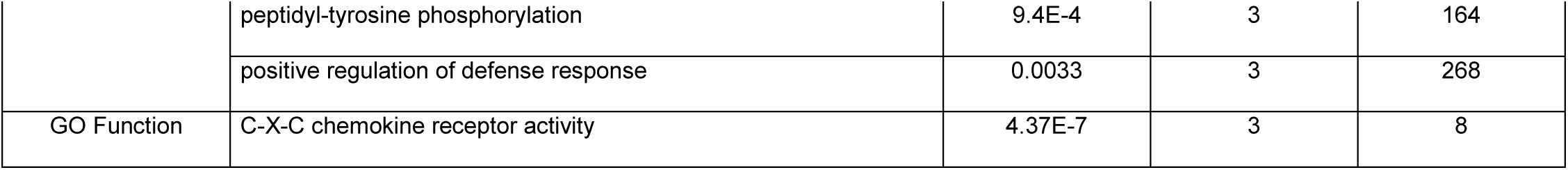
Cluster formation after functional enrichment analysis using STRING database. **A)** STRING interaction network. Minimum required interaction score is medium confidence (0.7). Node color: correlation with PC3. Parameters of Markov Cluster Algorithm: Granularity parameter (inflation value): 2.5. **(B)** Cluster C STRING functional enrichment. Similar terms were fused to reduce redundancy (Redundancy cut off= 0.25). The following figure supplements are available for figure 10: **Figure 11- figure supplement 1**. mRNA expression of particular genes at 1, 3, 5- and 10-days post infection.

In cluster C (**Figure 11**), the GO process “*positive regulation of interferon-gamma production*” (*Il12rb1, Il12rb2, Il23a, Il21, Il12b*) was overrepresented. In agreement, *Ifng* was upregulated at 10 dpi, suggesting that macrophage activation is occurring at 10 dpi and differentiation of CD4^+^ T cells is being promoted to the Th1 subset. This cytokine would probably be produced by NK cells and NKT cells (7), also by cells that have been activated by the innate immune response, specifically CD4^+^ T lymphocytes.

Cluster C **(Figure 11)** also shows the interaction among some genes (*Ccl7, Cxcr3, Ifng, Ccr7, Cxcr2, Cxcr5*) involved in the GO process: “*leukocyte chemotaxis*” GO.0030595. Nevertheless, the pathway with the lowest FDR in this cluster was “*cytokine-cytokine receptor interaction*”: mmu04060 (*Il12rb1, Il12rb2, Csf2, Ccl7, Il23a, Il21, Il21r, Cxcr3, Ifng, Ccr7, Cxcr2, Il12b, Cxcr5*) (**Figure 11 B**).

Strikingly, at 10 dpi, upregulation of *Pdcd1lg1* and *Pdcd1lg2* has been stopped, but there is remarkable upregulation of its ligand *Pdcd1*. Other regulatory markers were upregulated (*Lag3, Tim3*) at 10 dpi **(Figure 8, Figure 11- figure supplement 1)**.

## DISCUSSION

The most widely used approach for the study of differential gene expression under different experimental conditions is RNA-seq, but the results generally require confirmation by qPCR (1,13,25,26). In contrast, the development of high throughput Real-Time qPCR platforms allows large-scale gene-expression analysis of large collections of genes that can be used to identify global patterns (11). Our study involved High throughput Real-Time qPCR in livers of *L. infantum* infected (and non-infected, as controls) BALB/c mice very early after inoculation (1, 3, 5, 10-dpi). A multivariate statistical analysis was used to streamline this massive collection of transcriptomic data. Global PCA analysis revealed infected mice at 1- and 10 days post infection form two statistically significant independent groups, as a response to the expression of two separate sets of genes: those contributing to PC2 are overexpressed at 1 dpi (except *Rxra*) and those contributing to PC3 are overexpressed at 10 dpi. Therefore, we decided to base the analysis of the particular gene-expression signatures observed at those timepoints on the investigation of the molecular pathways and biological processes overrepresented in these gene sets.

The differentiation of those particular groups of mice was explained in the enrichment analysis (**Figure 3**). Our study demonstrated that at 1 dpi a variety of gene sets were enriched, that correspond to the upregulation of several genes involved in different processes in response to IFN-γ, mediators at membrane level, but also in cell chemotaxis. However, at 10 dpi the gene signature was characterized as a more targeted response with the upregulation of genes involved in *interleukin-12 signaling* pathway for the production of IFN-γ.

As early as 24 hours after parasite inoculation in mice, results of enrichment analysis and leading edge analysis (**Figure 5 C**), demonstrated that *Cxcl10* (IFN-γ Inducible Protein 10, IP-10) and *Cxcl9* (Monokine Induced by IFN-γ, MIG) are the most important markers in our study. These two chemokines were strongly upregulated at 1 dpi (**Figure 11- figure supplement 1**) and they are critical in both innate and adaptive immune responses to *Leishmania* infection (27). CXCL10 has been studied as a treatment in BALB/c mice infected by *L. infantum*, since there is evidence that CXCL10-treatment reduces IL-10^+^ Treg cell populations (CD4^+^CD25^+^Foxp3^+^ and Tr1) and induces morphological changes in the spleen (28). This chemokine also induces a reduction in parasite burden in the spleen and a decrease in IL-10 and TGF-β production in *L. infantum*-infected BALB/c mice (29).

At this timepoint, the upregulation of *Cxcl10* and *Cxcl9* are chemotactic signals to NK cells that would be generating IFN-γ at this timepoint, in agreement with the high enrichment of HALLMARK_INTERFERON_GAMMA_RESPONSE, and the upregulation of some genes as response to IFN-γ (**Figure 4 A**). It has been demonstrated that NK and NKT cells are sources of hepatic IFN-γ and this in turn is the responsible for the accumulation of CXCL10 mRNA at 1 dpi of hepatic *L. donovani* infection in C57BL/6 mice (19).

Once parasites are inoculated in the mouse and Kupffer macrophages uptake the parasites, one important cell population that participates at this early stage are natural killer T (NKT) cells (30,31) and particularly iNKT cells (31). The strong upregulation of *Cd69* at 1 dpi, an iNKT cell marker, suggests a key role for this cell type at this timepoint. Other genes correlated with PC2, such as *Cxcr4, Ccr4* and *Ccr5* are also expressed in NK cells (32) (**Figure 7**). These chemokine receptors and some other chemokines were also found in “*chemokine-mediated signaling pathway*” (**Figure 6B**), that involves some genes previously mentioned (*Cxcl10, Cxcl9)* and also *Ccl2* which is chemoattractant of macrophages, monocytes, NK cells and other CCR2-expressing leukocytes. CCL2 has been shown to play an important role in early immunity against cutaneous leishmaniasis (27,33).

These markers are probably secreted by the nascent granulomas (a nucleus of parasitized Kupffer cells surrounded by other immune cells) that start assembling once *Leishmania* parasites are transferred to the liver after inoculation (34). Those structures are crucial to activate the parasite-killing abilities of Kupffer cells (34), and secrete chemokines and cytokines that recruit immune cells, including monocytes, neutrophils and invariant natural T killer (iNKT) cells (6,7).

Additionally, at 1 dpi the upregulation of some genes involved in eicosanoids metabolism (*Alox12, Alox5, Pla2g4a)*, would contribute to the inflammatory process that exerts the infection in the liver to the formation of granuloma (31).

The upregulation of genes involved in *“arachidonic acid metabolism pathway”* (*Ptgds2, Ptges*) was also evident at this timepoint. PTGES is an arachidonic acid metabolite produced by immune cells, that has been recognized to support establishment of infection in macrophages, and hamsters infected by *L. donovani* (23).

The inflammatory chemokine response in Kupffer cells is driven by upstream signalling by Toll-like receptors (35,36). In our experiments, *Tlr2* and *Tlr4* were correlated with PC2. *Tlr2* has been reported as potential therapeutic target in visceral leishmaniasis (37). In livers of susceptible but self-curing C57BL/6 mice, *L. donovani* infection enhanced Toll-like receptor 4 (TLR4) and TLR2 gene expression. In TLR2(-/-) mice, control of liver infection, parasite killing, and granuloma assembly has been accelerated and chemotherapy’s efficacy enhanced (37).

The upregulation of some genes that were correlated to PC2, demonstrated the importance at 1 dpi, of some mediators at membrane level: *Tnfrsf18* (*Gitr*), *Itgam* (*Cd11b*), *Itgal* (*Cd11a*), *Itgb2, Icam1*. GITR has been studied as one candidate for immunomodulatory strategies (38). Faleiro et al., demonstrated anti-GITR mAb combined with anti-IL-10 as treatment in C57BL/6 mice infected with *L. donovani*, improved anti-parasitic immunity when used with sub-optimal doses of an anti-parasitic drug (38). However, it has been demonstrated with VL patient samples that targeting GITR alone in comparison with IL-10 signaling blockade, does not improve anti-parasitic immune responses, even with drug treatment cover.

ITGAM (CD11B) in combination with ITGB2 (CD18) form a leukocyte-specific integrin or macrophage receptor 1 (Mac-1). This receptor has been considered the predominant complement receptor responsible for the phagocytosis of complement-opsonized metacyclic promastigotes in the infection by *L. major* in human monocyte-derived macrophages (39).

ITGAL (CD11A), also correlated with PC2, in combination with ITGB2 form LFA-1, which has a central role in leukocyte intercellular adhesion through interactions with its ligands, ICAMs 1-3 (intercellular adhesion molecules 1 through 3), and also functions in lymphocyte costimulatory signaling (40). ITGB2, essential to form Mac1 and LFA-1, was upregulated at 1 dpi. *Icam1* was also upregulated at this timepoint. All these genes were correlated with PC2.

At 1 dpi, the upregulation of genes involved in “negative regulation of interferon-gamma production” (*Pdcd1lg1, Pdcd1lg2* and *Gata3*, (**Figure 6, Figure 8**)) could be a response to counterbalance the upregulation of Th1 markers and therefore restrain the response to IFNG at this timepoint. The interactions PDCD1-PD1L1 or PDCD1-PD2L2 are well known to inhibit T cell effector functions including IFN-γ production and proliferation, and the blockade of PD1L1 (41) and/or PD1L2 (12) contribute to the decrease of parasitic load.

The negative correlation of *Rxra* with PC2, is a consequence of its downregulation in 1 dpi-infected mice group. This gene has been reported like a central hub in the intracellular pathogen survival (1), is downregulated in KC in response to inflammation, but is aberrantly maintained in infected cells. It has been demonstrated that pharmacological manipulation of RXRa activity perturbs the transcriptomic network of infected KCs, leading to enhanced leishmanicidal activity (1). Our results suggest that, at 1 dpi, the downregulation of *Rxra*, is due to the role of KC in the inflammatory process at this timepoint. Later on the experiment, the consolidation of the infection in KC cells, may be key for reversing the downregulation of this gene in the liver.

All this information shows that 1 dpi, the differential pattern of gene expression in infected mice is related to inflammatory signals, triggered by chemokine response, eicosanoid metabolism and prostaglandin biosynthesis. This is the first step in the activation of innate immunity by inducing production of proinflammatory cytokines to culminate in the release of reactive oxygen species (ROS) and NO produced by macrophages (42,43).

At 10 days post infection, parasite load increased significantly (more than two orders of magnitude) compared to 1- and 3-days post infection. The statistically significant difference in the parasite burden between 1- and 10-days post inoculation, in addition to the hepatomegaly observed in 10 dpi-infected mice, supports the divergent gene signature that differentiate this two groups of mice. This is consistent with the full maturation of granuloma by 2-4 weeks after infection (34). Despite the fact that infection was clearly stablished as early as 1 dpi in the liver, inflammation was only evident a few days later, reflecting the time needed for the clinical signs to appear.

The gene signature at 10 dpi is characterized by the production of IFN-γ, after interleukin 12 stimulation, due to the upregulation of *I12b, Il12rb1* e *Il12rb2* (44). One of the main cytokines involved in the production of IFNG is IL12 which induce development of Th1 cells (45). We found upregulation of *Il12b* and its receptors *Il12rb1, Il12rb2*, all of them included in cluster C (**Figure 11 B**). Meanwhile, IL23 induce the differentiation of Th17 cells that, in turn, produce IL17, IL17F, IL6, and TNFα, but not IFNG and IL4 (reviewed in (46)). IL23 and IL12 have different effects (47), however those cytokines share the p40 subunit (IL12B). The receptor for IL23 is composed by IL12RB1 and IL23R and is important for the development of Th17 cell lineages (reviewed in (46)). Also, since IL12RB1, along with IL12RB2, are receptors for IL12, those genes are important for the Th1 lineage differentiation.

Despite the overrepresentation of “*IL17 signaling pathway*” (*Il12rb1, Il23a, Icosl, Il12b*), and the upregulation of some important mediators in the differentiation of Th17 lineage (*Il23a* and *Tbet*), the absence of the expression of *Il23r* (which in turn could be inhibited by IFNG and STAT1 (48)), would certainly inhibit the differentiation of this lineage. Down-regulation of *Il6* (**Figure 11- figure supplement 1**) also supports this idea, because this gene, along with TGFB are necessary in the differentiation of pathogenic Th17 cells from naive T cells (49).

Another sign of inhibition of inflammatory Th17 lineage development at 10 dpi is the upregulation of *Il27 (p28)* and *Ebi3* (GO process “*positive regulation of interferon-gamma production*”). Both form IL27, whose receptor IL27R is composed of gp130 (IL6ST) and IL27RA (50). All these mediators were upregulated in our data except *Il6st* (**Figure 11- figure supplement 1**), that was expressed at similar levels in infected and control mice. The interaction IL27-IL27 receptor is an early signal for the induction of Th1 response (51) and could inhibit inflammatory Th17 lineage development (reviewed in (52), (53)). Both facts are consistent with our results. Additionally, IL27 synergize with IL12 the production of IFNG by CD4, CD8 T cells and NKT cells (54,55). In the interaction network we found that all those genes are involved in IFNG production **(Figure 11)**.

At 10 dpi, the upregulation of *Cxcr3* (Th1-specific chemokine receptor (reviewed in (56)) and *Ccl7* (chemoattractant of monocytes, eosinophils and basophils (57)) suggest there is active signaling of Th1/M1 response cells at the site of the infection at 10 dpi. These signals for leukocyte recruitment trigger granuloma formation in the liver to the expected full maturation by 2–4 weeks after infection (58). The upregulation of TNF (Figure 11- figure supplement 1) also support this, since this marker plays a crucial role in coordinating the assembly and maturation of granulomas (reviewed in (8,30)).

Some regulatory markers were upregulated at 10 dpi (*Pdcd1, Lag3, Tim3*). These genes have been involved in T cell exhaustion not only in chronic viral infections (reviewed in (59)), but also in protozoan diseases (reviewed in (60)). Although PDCD1, TIM3 and LAG3 can participate synergistically, in non-redundant signaling pathways towards T cells exhaustion (61), the actual role of these signals in our model is unclear. Despite the upregulation of these regulatory markers at 10 dpi, the upregulation of IL-12 signaling gene set would stimulate the positive regulation of interferon-gamma production. This is expected to control the parasite replication in posterior days in the liver.

The information derived from the expression profile of genes correlated with PC3 suggest that the gene signature at this timepoint is more clearly defined towards activation of Th1/M1 responses through interleukin-12 signaling and positive regulation of interferon-gamma production, as well as inhibition of inflammatory Th17 lineage development. Despite the upregulation of T cell exhaustion genes at 10 dpi, their role is unclear (**Figure 11 A, figure 11 B**).

Numerous scientists have used transcriptional analysis as a tool to detect changes in complex biological scenarios, such as infection or candidate vaccine testing. Most of those analysis either lack in precision due to the limited number of genes analyzed, or, on the contrary, use RNA-seq methodologies that provide a plethora of different data, sometimes difficult to relate to the topic of interest, and that needs to be confirmed using RTqPCR. Our global multivariate statistical approach applied to a large-scale gene expression experiment allowed to streamline a large collection of RT-PCR gene expression data in order to decipher the immunological mechanisms underlying early *L. infantum* infection in livers of BALB/c mice. This novel approach simplifies the characterization of the most relevant changes at the transcriptome level and the connection with other data such as parasite burden and weight of organ, providing a more accurate insight on the changes over time during infection. The evaluation of transcriptional changes in the early *L. infantum* infection can also lead us to identify possible candidates to biomarkers and immunomodulatory strategies. One of this candidate molecules is *Rxra*, that shows a particular pattern of expression, inversely proportional to the expression of key Th1/M1 markers like *Cxcl9* and *Cxcl10* at 1 dpi. Given the switch in its expression pattern during infection and its central role in the transcriptomic network of infected KCs, *Rxra* should be the focus of future studies to investigate the role of this gene as a potential target to control *L. infantum* infection.

## MATERIALS AND METHODS

### Biological samples

Experiments involving animals were conducted in accordance to both European (2010/63/UE) and Spanish legislation (Law 53/2013), after approval by the Committee for Research Ethics and Animal Welfare (CEIBA) of the University of La Laguna (Permission code: CEIBA2015-0168).

Forty-eight female wild-type BALB/c mice were obtained from the breeding facilities of the Charles River Laboratories, (France) and were maintained under specific pathogen-free conditions at ULL. Half of these animals (n=24) were randomly assigned to control group and the other half (n=24) were infected by inoculation in the coccygeal vein of 10^6^ promastigotes of *L. infantum* (JPC strain, MCAN/ES/98/LLM-724) in stationary growth phase on day 0 of the experiment. *L. infantum* was maintained in vivo by serial murine passages. Prior to infection, amplification of amastigote-derived promastigotes, with less than 3 passages in vitro, was carried out by culture in RPMI medium (Gibco BRL), supplemented with 20% inactivated fetal calf serum (SBFI), 100 ug/ml streptomycin (Sigma-Aldrich, St. Louis, USA) and 100 U/ml of penicillin (Biochrom AG, Berlin, Germany) at 26°C until reaching stationary phase.

At 1, 3, 5, 10 days post-infection (dpi), 6 infected mice and 6 control mice (n=12 each timepoint) were euthanized by cervical dislocation to extract the liver and proceed to the determination of liver weight and parasitic load (62). Samples were immediately stored in RNA later at −80°C (Sigma-Aldrich, St. Louis, USA) for nucleic acid preservation and further mRNA extraction.

### RNA isolation and quantification

Preserved liver tissue (7-10 mg) was homogenized with TRI Reagent (Sigma-Aldrich) in Lysing Matrix D columns (MP Biomedicals) and FastPrep® System (ProScientific). mRNA enrichment was performed using RNeasy Mini Kit (Qiagen), following manufacturer’s instructions. *DeNovix DS-11 Spectrophotometer* (*DeNovix*) was used to quantify and evaluate RNA purity. Only samples with OD_260/280_ ratios between 2-2.2 and ratio OD_260/230_ between 1.8-2.2 were included in this study. The integrity of the RNA was determined using 2100 Bioanalyzer (Agilent Technologies) and RNA 6000 Nano Kit chips (Agilent Technologies). RIN (RNA integrity number) was >7 for all RNA samples included in this study. Additionally, agarose gel electrophoresis was performed to all RNA samples for further verification of integrity and quality.

### Retrotranscription and High-Throughput Real-Time Quantitative PCR (RT-qPCR)

*High Capacity cDNA Reverse Transcription kit* (Thermo Fisher) was used to perform the retrotranscription using a Veriti® 96-Well Fast Thermal Cycler thermocycler (Thermo Fisher). Real Time qPCR was performed using OpenArray® plates with TaqMan probes (Thermo Fisher) for the amplification of 223 genes that were selected and related to innate and adaptive immune response, prostaglandin synthesis, lipid metabolism, C-type lectin receptors and MAPK signaling pathway. All primers and probes were commercially designed by Thermo Fisher Scientific (**Supplementary file 1**). Each plate allowed performing 3072 amplification reactions. The plate was organized in 48 subarrays, each with 64 nanopockets. PCR mixture was prepared following the manufacturer’s instructions and plates were loaded using the Accufill TM System (Thermo Fisher Scientific) Each amplification reaction was performed in a volume of 33 nl. The thermal cycle and fluorescence detection were performed with the QuantStudio 12K Flex Real-Time PCR System (Thermo Fisher Scientific) following manufacturer’s recommendations. Each subarray contained the primers together with a probe labeled with FAM at its 5 ‘end. Each sample was amplified in triplicate. *Cq* values produced by this platform are already corrected for the efficiency of the amplification (11,63) and candidate reference genes.

### Data pre-processing and processing

Data were exported from QuantStudioTM 12K Flex Real-Time PCR System software to MS Excel. The arithmetic average quantitative cycle (Cq) was used for data analysis. *GenEX (MultiD)* software was used for data preprocessing and normalization. Reference genes selection for this dataset (63) was performed in *GenEx* using two different methods: geNorm (setting M-value lower than 0.5) and NormFinder, revealing two genes that showed the most stable expression: *Lta4h, P38mapk*. Normalization with these genes was performed to obtain NRQ values (Normalized Relative Quantities).

### Statistical analysis

Mean and its standard error (SEM) was calculated for each parameter measured. For all the analyses, outliers were assessed by inspection of boxplots and studentized residuals, data were checked for normal distribution by the Kolmogorov– Smirnoff test or Shapiro Wilk test, as appropriate, and homogeneity of the variance by means of the Levene test.

The differences in parasite burden between time points were determined by a one-way Analysis of Variance (ANOVA) and a two-way ANOVA was used to compare the weight of liver between infected and control mice at each time point. Tukey’s test and pairwise comparisons with Bonferroni adjustment as post hoc comparison techniques were carried out to compare the means from different groups.

NRQ values were used to perform a Principal Component Analysis (PCA), an unsupervised multivariate method, which provides a first approximation to the statistical analysis of expression profiling data with the aim of generate several hypotheses of interest. In the PCA, the correlation between variables was assessed by inspection of the correlation matrix, the overall Kaiser-Meyer-Olkin (KMO) measure and the Bartlett’s test of sphericity, indicating that the data were likely factorizable.

Unsupervised hierarchal clustering of the NRQ values was applied and based on Euclidean distance and pairwise average-linkage was performed in Cytoscape 3.8.0 (http://www.cytoscape.org/) (64) using the plugin clusterMaker (65).

Next step was a confirmatory study to test the hypothesis; differences in Principal Component 2 (PC2) and Principal Component 3 (PC3) scores were determined by a two-way ANOVA to study the combined effects of both factors, time post infection (1-,3-,5- and 10-days post infection) and condition (infected or control), as well as their interactions, followed by Tukey’s post-hoc tests and pairwise comparisons with Bonferroni adjustment, as appropriate.

Descriptive analysis, the evaluation of assumptions, ANOVA’s and Principal Component Analysis were carried out using IBM^®^ SPSS^®^ version 26 (IBM Corporation, Armonk, NY) statistical software.

NRQ values were also used for individual gene representation. Differences in NRQ values between non-infected (control) and infected animals were analyzed by two tail Mann-Whitney test or two tail unpaired t-test, as appropriate, using GraphPad Prism version 8, GraphPad Software, San Diego California USA (www.graphpad.com).

### Enrichment analysis

To analyze the potential biological processes and pathways regarding to PC2- and PC3-correlated genes, we used the software Gene Set Enrichment Analysis (GSEA) version 4.0.3. This is a computational method that determines whether an *a priori* defined set of genes shows statistically significant differences between two biological states (e.g. phenotypes) (17). Metric for ranking genes was tTest ratio. 1000 gene permutations were used to generate a null distribution for ES (Enrichment score), then each pathway will attain a normalization enrichment score (NES). We performed the enrichment analysis using gene sets from Gene Ontology (GO) biological process excluding annotations that have evidence code IEA (inferred from electronic annotation), ND (no biological data available), and RCA (inferred from reviewed computational analysis) and including all pathway resources (Reactome, Panther, NetPath, NCI Nature, MSigdb, IOB, WikiPathways).

With enrichment results, the Cytoscape *3*.*8*.*0*. software and the collection of plugins that are part of the EnrichmentMap pipeline workflow (66) were used to create enrichment maps in order to integrate the results from GSEA. The enrichment map was generated using only the gene-sets passing the thresholds: nominal *p*-value < 0.001, *q*-value < 0.25. The overlap coefficient was set to 0.35. The plugin AutoAnnote was used to perform Markov-based flow simulation to determine clusters (Markov Cluster Algorithm).

Leading edge analysis was performed in GSEA to elucidate key genes associated with the Top 5 enriched gene sets.

### PPI network

We used STRING database (STRING, http://string-db.org) (24) to visualize the interactive relationships of the PC2- and PC3-correlated genes. Confidence score > 0.7 was considered as significant. Then, the Cytoscape software was used for constructing and visualizing PPI networks.

## ACKNOWLEDGMENTS

We would like to thank Dr. M.C. Lopez and Dr. MC. Thomas (IPBLN-CSIC, Spain) for helpful discussions after reading this manuscript.

## ADDITIONAL INFORMATION

### Author Contributions

Génesis Palacios Data curation, Software, Formal analysis, Validation, Investigation, Visualization, Methodology, Writing—original draft; Raquel Diaz-Solano, Methodology, Validation; Basilio Valladares, Supervision, Funding acquisition, Writing—review and editing; Roberto Dorta-Guerra, Data curation, Software, Validation, Visualization, Writing— review and editing; Emma Carmelo, Conceptualization, Resources, Supervision, Funding acquisition, Methodology, Formal analysis, Validation, Investigation, Writing—original draft, Project administration, Writing—review and editing.

### Data availability

Data used in this study have been deposited in GEO under accession code GSE159195.

