## Supplemental material for "Integrated multivariate analysis of transcriptomic data reveals immunological mechanisms in mice after *Leishmania infantum* infection"

[1] Instituto Universitario de Enfermedades Tropicales y Salud Pública de Canarias, Universidad de la Laguna, La Laguna, Islas Canarias, Spain.[2] Departamento de Obstetricia y Ginecología, Pediatría, Medicina Preventiva y Salud Pública, Toxicología, Medicina Legal y Forense y Parasitología. [3] Departamento de Matemáticas, Estadística e Investigación Operativa, Facultad de Ciencias, Universidad de La Laguna, La Laguna, Spain. [4] Red de Investigación Colaborativa en Enfermedades Tropicales (RICET).

### SUPPLEMENTAL MATERIAL

**Supplementary file 1.** List of TaqMan assays used for RT-qPCR analysis using QuantStudioTM 12K Flex Real-Time PCR System.

PANEL 1.

| N° | Assay ID | GEN | NOMBRE DEL GEN | GRUPO |
|---|---|---|---|---|
| 1 | Mm00437762_m1 | B2m | beta-2 microglobulin | Endogenous gene expression |
| 2 | Mm00446968_m1 | Hprt | hypoxanthine guanine phosphoribosyl transferase | |
| 3 | Mm00435617_m1 | Pgk1 | phosphoglycerate kinase 1 | |
| 4 | Mm01277042_m1 | Tbp | TATA box binding protein | |
| 5 | Mm01201237_m1 | Ubc | ubiquitin C | |
| 6 | Mm01722325_m1 | Ywhaz | tyrosine 3-monooxygenase/tryptophan 5-monooxygenase activation protein, zeta polypeptide | |
| 7 | Mm00599890_m1 | Ifngr1 | interferon gamma receptor 1 | Cytokines and cytokine receptors |
| 8 | Mm00492626_m1 | Ifngr2 | interferon gamma receptor 2 | |
| 9 | Mm01168134_m1 | Ifng | interferon gamma | |
| 10 | Mm01178820_m1 | Tgfb1 | transforming growth factor, beta 1 | |
| 11 | Mm00436955_m1 | Tgfb2 | transforming growth factor, beta 2 | |
| 12 | Mm00436964_m1 | Tgfbr1 | transforming growth factor, beta receptor I | |
| 13 | Mm00436977_m1 | Tgfbr2 | transforming growth factor, beta receptor II | |
| 14 | Mm00443258_m1 | Tnf | tumor necrosis factor | |
| 15 | Mm00441875_m1 | Tnfrsf1a | tumor necrosis factor receptor superfamily, member 1a | |
| 16 | Mm00441889_m1 | Tnfrsf1b | tumor necrosis factor receptor superfamily, member 1b | |
| 17 | Mm01288580_m1 | Irf1 | interferon regulatory factor 1 | |
| 18 | Mm00496 7_m1 | Irf5 | interferon regulatory factor 5 | |
| 19 | Mm00516788_m1 | Irf7 | interferon regulatory factor 7 | |
| 20 | Mm01224532_m1 | Irg1 | immunoresponsive gene 1 | |
| 21 | Mm00518984_m1 | Il23a | interleukin 23, alpha subunit p19 | Interleukines and interleukin receptors |
| 22 | Mm00434237_m1 | Il1r1 | interleukin 1 receptor, type I | |
| 23 | Mm00439614_m1 | Il10 | interleukin 10 | |

| | | | |
|---|---|---|---|
| 24 | Mm00434157_m1 | Il10rb | interleukin 10 receptor, beta |
| 25 | Mm00434169_m1 | Il12a | interleukin 12a |
| 26 | Mm00434174_m1 | Il12b | interleukin 12b |
| 27 | Mm00434189_m1 | Il12rb1 | interleukin 12 receptor, beta 1 |
| 28 | Mm00434200_m1 | Il12rb2 | interleukin 12 receptor, beta 2 |
| 29 | Mm00434204_m1 | Il13 | interleukin 13 |
| 30 | Mm00439618_m1 | Il17a | interleukin 17A |
| 31 | Mm00521423_m1 | Il17f | interleukin 17F |
| 32 | Mm00434214_m1 | Il17ra | interleukin 17 receptor A |
| 33 | Mm00434226_m1 | Il18 | interleukin 18 |
| 34 | Mm00456733_m1 | Il18bp | interleukin 18 binding protein |
| 35 | Mm00439620_m1 | Il1a | interleukin 1 alpha |
| 36 | Mm00434228_m1 | Il1b | interleukin 1 beta |
| 37 | Mm00446186_m1 | Il1rn | interleukin 1 receptor antagonist |
| 38 | Mm00434256_m1 | Il2 | interleukin 2 |
| 39 | Mm00517640_m1 | Il21 | interleukin 21 |
| 40 | Mm00600317_m1 | Il21r | interleukin 21 receptor |
| 41 | Mm01192969_m1 | Il22ra2 | interleukin 22 receptor, alpha 2 |
| 42 | Mm00519943_m1 | Il23r | interleukin 23 receptor |
| 43 | Mm00461162_m1 | Il27 | interleukin 27 |
| 44 | Mm00497259_m1 | Il27ra | interleukin 27 receptor, alpha |
| 45 | Mm00442885_m1 | Il2rg | interleukin 2 receptor, gamma chain |
| 46 | Mm00445259_m1 | Il4 | interleukin 4 |
| 47 | Mm00439646_m1 | Il5 | interleukin 5 |
| 48 | Mm00434284_m1 | Il5ra | interleukin 5 receptor, alpha |
| 49 | Mm00446190_m1 | Il6 | interleukin 6 |
| 50 | Mm00439653_m1 | Il6ra | interleukin 6 receptor, alpha |
| 51 | Mm00439665_m1 | Il6st | interleukin 6 signal transducer |
| 52 | Mm01290062_m1 | Csf2 | colony stimulating factor 2 (granulocyte-macrophage) |
| 53 | Mm00655745_m1 | Csf2rb | colony stimulating factor 2 receptor, beta, low-affinity (granulocyte-macrophage) |
| 54 | Mm00469294_m1 | Ebi3 | Epstein-Barr virus induced gene 3 |

| | | | | |
|---|---|---|---|---|
| 55 | Mm00475988_m1 | Arg1 | arginase, liver | |
| 56 | Mm00475162_m1 | Foxp3 | forkhead box P3 | |
| 57 | Mm00440338_m1 | Myd88 | myeloid differentiation primary response gene 88 | Enzymes, prostaglandins and adaptor proteins |
| 58 | Mm00440502_m1 | Nos2 | nitric oxide synthase 2, inducible | |
| 59 | Mm00479246_m1 | Nox4 | NADPH oxidase 4 | |
| 60 | Mm00478374_m1 | Ptgs2 | prostaglandin-endoperoxide synthase 2 | |
| 61 | Mm00439531_m1 | Stat1 | signal transducer and activator of transcription 1 | |
| 62 | Mm01219775_m1 | Stat3 | signal transducer and activator of transcription 3 | |
| 63 | Mm00448890_m1 | Stat4 | signal transducer and activator of transcription 4 | |
| 64 | Mm01160477_m1 | Stat6 | signal transducer and activator of transcription 6 | |
| 65 | Mm00477633_m1 | Bcl6 | B cell leukemia/lymphoma 6 | |
| 66 | Mm00492590_m1 | Ido1 | indoleamine 2,3-dioxygenase 1 | |
| 67 | Mm00500554_m1 | Mmp12 | matrix metallopeptidase 12 | |
| 68 | Mm00485054_m1 | Mmp14 | matrix metallopeptidase 14 (membrane-inserted) | |
| 69 | Mm00476361_m1 | Nfkb1 | nuclear factor of kappa light polypeptide gene enhancer in B cells 1, p105 | |
| 70 | Mm00479807_m1 | Nfkb2 | nuclear factor of kappa light polypeptide gene enhancer in B cells 2, p49/p100 | |
| 71 | Mm00450960_m1 | Tbx21 | T-box 21 | |
| 72 | Mm00441891_m1 | Cd40 | CD40 antigen | Costimulatory and cell adhesion molecules |
| 73 | Mm00441911_m1 | Cd40lg | CD40 ligand | |
| 74 | Mm00711660_m1 | Cd80 | CD80 antigen | |
| 75 | Mm00444543_m1 | Cd86 | CD86 antigen | |
| 76 | Mm00486849_m1 | Ctla4 | cytotoxic T-lymphocyte-associated protein 4 | |
| 77 | Mm03048248_m1 | Cd274 | CD274 antigen | |
| 78 | Mm00516023_m1 | Icam1 | intercellular adhesion molecule 1 | |
| 79 | Mm00494862_m1 | Icam2 | intercellular adhesion molecule 2 | |
| 80 | Mm00497600_m1 | Icos | inducible T cell co-stimulator | |
| 81 | Mm00497237_m1 | Icosl | icos ligand | |
| 82 | Mm00434513_m1 | Itgb2 | integrin beta 2 | |
| 83 | Mm00486868_m1 | Cd83 | CD83 antigen | |
| 84 | Mm00493071_m1 | Lag3 | lymphocyte-activation gene 3 | |
| 85 | Mm01285676_m1 | Pdcd1 | programmed cell death 1 | |

| | | | | |
|---|---|---|---|---|
| 86 | Mm00454540_m1 | Havcr2 | hepatitis A virus cellular receptor 2 | |
| 87 | Mm00441242_m1 | Ccl2 | chemokine (C-C motif) ligand 2 | |
| 88 | Mm00441258_m1 | Ccl3 | chemokine (C-C motif) ligand 3 | |
| 89 | Mm00443111_m1 | Ccl4 | chemokine (C-C motif) ligand 4 | |
| 90 | Mm01302427_m1 | Ccl5 | chemokine (C-C motif) ligand 5 | |
| 91 | Mm00443113_m1 | Ccl7 | chemokine (C-C motif) ligand 7 | |
| 92 | Mm01216147_m1 | Ccr1 | chemokine (C-C motif) receptor 1 | |
| 93 | Mm01216173_m1 | Ccr2 | chemokine (C-C motif) receptor 2 | |
| 94 | Mm01216171_m1 | Ccr5 | chemokine (C-C motif) receptor 5 | |
| 95 | Mm01301785_m1 | Ccr7 | chemokine (C-C motif) receptor 7 | |
| 96 | Mm04207460_m1 | Cxcl1 | chemokine (C-X-C motif) ligand 1 | |
| 97 | Mm00445235_m1 | Cxcl10 | chemokine (C-X-C motif) ligand 10 | Chemokines and chemokine receptors |
| 98 | Mm00436450_m1 | Cxcl2 | chemokine (C-X-C motif) ligand 2 | |
| 99 | Mm00434946_m1 | Cxcl9 | chemokine (C-X-C motif) ligand 9 | |
| 100 | Mm00438258_m1 | Cxcr2 | chemokine (C-X-C motif) receptor 2 | |
| 101 | Mm00438259_m1 | Cxcr3 | chemokine (C-X-C motif) receptor 3 | |
| 102 | Mm00434772_m1 | Xcl1 | chemokine (C motif) ligand 1 | |
| 103 | Mm00441260_m1 | Ccl9 | chemokine (C-C motif) ligand 9 | |
| 104 | Mm00444533_m1 | Cxcl13 | chemokine (C-X-C motif) ligand 13 | |
| 105 | Mm00469712_m1 | Cxcl16 | chemokine (C-X-C motif) ligand 16 | |
| 106 | Mm01701838_m1 | Cxcl3 | chemokine (C-X-C motif) ligand 3 | |
| 107 | Mm00432086_m1 | Cxcr5 | chemokine (C-X-C motif) receptor 5 | |
| 108 | Mm00442346_m1 | Tlr2 | toll-like receptor 2 | |
| 109 | Mm01207404_m1 | Tlr3 | toll-like receptor 3 | |
| 110 | Mm00445273_m1 | Tlr4 | toll-like receptor 4 | Toll-like receptors |
| 111 | Mm00446590_m1 | Tlr7 | toll-like receptor 7 | |
| 112 | Mm00446193_m1 | Tlr9 | toll-like receptor 9 | |

PANEL 2.

| N° | Assay ID | GEN | NOMBRE DEL GEN | GRUPO |
|---|---|---|---|---|
| 1 | Mm00437762_m1 | B2m | beta-2 microglobulin | Endogenous gene expression |
| 2 | Mm00446968_m1 | Hprt | hypoxanthine guanine phosphoribosyl transferase | |
| 3 | Mm01201237_m1 | Ubc | ubiquitin C | |
| 4 | Mm01331626_m1 | Akt1 | thymoma viral proto-oncogene 1 | MAPK signaling pathway |
| 5 | Mm01173094_m1 | Akt2 | thymoma viral proto-oncogene 2 | |
| 6 | Mm00442194_m1 | Akt3 | thymoma viral proto-oncogene 3 | |
| 7 | Mm01973540_g1 | Mapk3, Erk1 | mitogen-activated protein kinase 3 | |
| 8 | Mm00442479_m1 | Mapk1, Erk2 | mitogen-activated protein kinase 1 | |
| 9 | Mm00489514_m1 | Mapk8, Jnk | mitogen-activated protein kinase 8 | |
| 10 | Mm00444968_m1 | Mtor | mechanistic target of rapamycin (serine/threonine kinase) | |
| 11 | Mm01301009_m1 | Mapk14, P38mapk | mitogen-activated protein kinase 14 | |
| 12 | Mm01282781_m1 | Pik3r1, Pi3k | phosphatidylinositol 3-kinase, regulatory subunit, polypeptide 1 (p85 alpha) | |
| 13 | Mm00440940_m1 | Pparg | peroxisome proliferator activated receptor gamma | Lipid metabolism |
| 14 | Mm00443451_m1 | Nr1h3 | nuclear receptor subfamily 1, group H, member 3 | |
| 15 | Mm00447040_m1 | Pla2g4a | phospholipase A2, group IVA (cytosolic, calcium-dependent) | |
| 16 | Mm00447271_m1 | Ptgis | prostaglandin I2 (prostacyclin) synthase | |
| 17 | Mm00441185_m1 | Rxra | retinoid X receptor alpha | |
| 18 | Mm00436051_m1 | Ptger2, Ep | prostaglandin E receptor 2 (subtype EP2) | Protaglandine synthesis |
| 19 | Mm00436053_m1 | Ptger4 | prostaglandin E receptor 4 (subtype EP4) | |
| 20 | Mm00452105_m1 | Ptges | prostaglandin E synthase | |
| 21 | Mm00460181_m1 | Ptges2 | prostaglandin E synthase 2(Ptges2) | |
| 22 | Mm01731378_g1 | Ptges3 | prostaglandin E synthase 3 (cytosolic) | |
| 23 | Mm00477214_m1 | Ptgs1 | prostaglandin-endoperoxide synthase 1 | |
| 24 | Mm01199500_m1 | P2rx7 | purinergic receptor P2X, ligand-gated ion channel, 7 | |
| 25 | Mm00435472_m1 | P2ry2 | purinergic receptor P2Y, G-protein coupled 2 | |
| 26 | Mm00436055_m1 | Ptgfr | prostaglandin F receptor | |
| 27 | Mm00479846_m1 | Hpgds, Ptgds2 | intercellular adhesion molecule 1 | |
| 28 | Mm00482476_m1 | Ptgr1, Ltb4dh | prostaglandin reductase 1 | |

| | | | | |
|---|---|---|---|---|
| 29 | Mm00521839_m1 | Ltb4r1, Blt1 | leukotriene B4 receptor 1 | |
| 30 | Mm01182747_m1 | Alox5 | arachidonate 5-lipoxygenase | |
| 31 | Mm00507789_m1 | Alox15 | arachidonate 15-lipoxygenase | |
| 32 | Mm00521826_m1 | Lta4h | leukotriene A4 hydrolase | |
| 33 | Mm00545833_m1 | Alox12 | arachidonate 12-lipoxygenase | |
| 34 | Mm00839636_g1 | Cd68 | CD68 antigen | Cholesterol pathway |
| 35 | Mm00432403_m1 | Cd36 | CD36 antigen | |
| 36 | Mm00459972_m1 | Cd209d | CD209d antigen | C-Type Lectin Receptors (CLRs) |
| 37 | Mm01183349_m1 | Clec7a | C-type lectin domain family 7, member a | |
| 38 | Mm01329362_m1 | Mrc1 | mannose receptor, C type 1 | |
| 39 | Mm01183378_m1 | Cd69 | CD69 antigen | |
| 40 | Mm00495182_m1 | Klrd1, Cd94 | killer cell lectin-like receptor, subfamily D, member 1 | |
| 41 | Mm00435587_m1 | Pfkl | phosphofructokinase, liver, B-type | Carbohidrates synthesis |
| 42 | Mm00439344_m1 | Hk1 | hexokinase 2 | |
| 43 | Mm00443385_m1 | Hk2 | hematopoietic prostaglandin D synthase | |
| 44 | Mm00441480_m1 | Slc2a1 | solute carrier family 2 (facilitated glucose transporter), member 1 | |
| 45 | Mm00472712_m1 | Gys1 | granzyme B | |
| 46 | Mm00504650_m1 | Pfkfb3 | 6-phosphofructo-2-kinase/fructose-2,6-biphosphatase 3 | |
| 47 | Mm00554300_m1 | Pdk1 | pyruvate dehydrogenase kinase, isoenzyme 1 | |
| 48 | Mm01268229_m1 | Pgm2 | phosphoglucomutase 2 | |
| 49 | Mm01612132_g1 | Ldha | lactate dehydrogenase A | |
| 50 | Mm00833691_g1 | Tpi1 | triosephosphate isomerase 1 | |
| 51 | Mm00434151_m1 | Il10ra | interleukin 10 receptor, beta | Interleukines and interleukin receptors |
| 52 | Mm00446726_m1 | Il13ra1 | interleukin 13 receptor, alpha 2 | |
| 53 | Mm00515166_m1 | Il13ra2 | interleukin 15 | |
| 54 | Mm00434210_m1 | Il15 | interleukin 16 | |
| 55 | Mm00516039_m1 | Il16 | interleukin 17 receptor A | |
| 56 | Mm00515178_m1 | Il18r1 | interleukin 18 receptor accessory protein | |
| 57 | Mm00516053_m1 | Il18rap | interleukin 1 alpha | |
| 58 | Mm00663697_m1 | Il22ra1 | interleukin 23, alpha subunit p19 | |
| 59 | Mm01340213_m1 | Il27ra | interleukin 2 receptor, alpha chain | |
| 60 | Mm00434268_m1 | Il2rb | interleukin 2 receptor, gamma chain | |

| | | | | |
|---|---|---|---|---|
| 61 | Mm01275139_m1 | Il4ra | interleukin 5 receptor, alpha | |
| 62 | Mm00434295_m1 | Il7r, Cd127 | interleukin 9 receptor | |
| 63 | Mm00434313_m1 | Il9r | interferon regulatory factor 1 | |
| 64 | Mm00499822_m1 | Il25 | interleukin 25 | |
| 65 | Mm00434305_m1 | Il9 | interleukin 9 | |
| 66 | Mm00499822_m1 | Il2ra, Cd25 | interleukin 2 receptor, alpha chain | |
| 67 | Mm00444241_m1 | Il22 | interleukin 22 | |
| 68 | Mm00516136_m1 | Ccl17 | chemokine (C-C motif) ligand 17 | Chemokines and chemokine receptors |
| 69 | Mm00839966_g1 | Ccl19 | chemokine (C-C motif) ligand 19 | |
| 70 | Mm00436439_m1 | Ccl22 | chemokine (C-C motif) ligand 22 | |
| 71 | Mm00438271_m1 | Ccr4 | chemokine (C-C motif) receptor 4 | |
| 72 | Mm00444662_m1 | Cxcl11 | chemokine (C-X-C motif) ligand 11 | |
| 73 | Mm01292123_m1 | Cxcr4 | chemokine (C-X-C motif) receptor 4 | |
| 75 | Mm00472858_m1 | Cxcr6 | Epstein-Barr virus induced gene 3 | |
| 76 | Mm00441263_m1 | Cxcl15 | chemokine (C-X-C motif) ligand 15 | |
| 77 | Mm00490880_m1 | Stat2 | signal transducer and activator of transcription 2 | Enzymes and adaptor proteins |
| 78 | Mm00600614_m1 | Jak1 | Janus kinase 1 | |
| 79 | Mm01208489_m1 | Jak2 | Janus kinase 2 | |
| 80 | Mm00477631_m1 | Bcl2 | B cell leukemia/lymphoma 2 | |
| 81 | Mm00432050_m1 | Bax | BCL2-associated X protein | |
| 82 | Mm00437783_m1 | Bcl2l1 | BCL2-like 1 | |
| 83 | Mm00438861_m1 | Fadd | Fas (TNF receptor superfamily member 6) | |
| 84 | Mm00442834_m1 | Gzmb | hepatitis A virus cellular receptor 2 | |
| 85 | Mm00436979_m1 | Tgm2 | transglutaminase 2, C polypeptide | |
| 86 | Mm00445109_m1 | Retnla, Fizz1 | resistin like alpha | |
| 87 | Mm00448427_m1 | Ptpn1, Ptp1b | protein tyrosine phosphatase, non-receptor type 1 | |
| 88 | Mm01278617_m1 | Mki67, Ki67 | antigen identified by monoclonal antibody Ki 67 | |
| 89 | Mm00661498_m1 | CD57, B3gat1 | beta-1,3-glucuronyltransferase 1 (glucuronosyltransferase P) | |
| 90 | Mm00451734_m1 | Pdcd1lg2 | programmed cell death 1 ligand 2 | Costimulatory and cell adhesion molecules |
| 91 | Mm00599683_m1 | Cd3e | CD3 antigen, epsilon polypeptide | |
| 92 | Mm01182108_m1 | Cd8a | CD8 antigen, alpha chain | |
| 93 | Mm00488332_m1 | Siglec1, Cd169 | sialic acid binding Ig-like lectin 1, sialoadhesin | |
| 94 | Mm00434455_m1 | Itgam | integrin alpha M | |

| | | | | |
|---|---|---|---|---|
| 95 | Mm00483137_m1 | Cd28 | CD28 antigen | |
| 96 | Mm00801807_m1 | Itgal | integrin alpha M | |
| 97 | Mm00444461_m1 | Cd160 | CD160 antigen | |
| 98 | Mm01251919_m1 | Itgae, Cd103 | integrin alpha L | |
| 99 | Mm00442754_m1 | Cd4 | CD4 antigen(Cd4) | |
| 100 | Mm01149710_m1 | Ncam1, Cd56 | neural cell adhesion molecule 1 | |
| 101 | Mm00484683_m1 | Gata3 | glycogen synthase 1, muscle | |
| 102 | Mm00467257_m1 | Nfat5 | nuclear factor of activated T cells 5 | Transcription factors |
| 103 | Mm00468869_m1 | Hif1a | hexokinase 1 | |
| 104 | Mm00627599_m1 | Ucp2 | uncoupling protein 2 (mitochondrial, proton carrier) | Carrier proteins |
| 105 | Mm00437136_m1 | Tnfrsf18 | tumor necrosis factor receptor superfamily, member 18 | |
| 106 | Mm00616981_m1 | Btla | B and T lymphocyte associated | |
| 107 | Mm00438864_m1 | Fasl | forkhead box P3 | |
| 108 | Mm01204974_m1 | Fas | Fas ligand (TNF superfamily, member 6) | Cytokines and cytokine receptors |
| 109 | Mm00442039_m1 | Tnfrsf4 | tumor necrosis factor receptor superfamily, member 4 | |
| 110 | Mm00440228_gH | Lta | lymphotoxin A | |
| 111 | Mm01342740_g1 | Socs1 | suppressor of cytokine signaling 1 | |
| 112 | Mm01249143_g1 | Socs3 | suppressor of cytokine signaling 3 | |

### Figure 2- figure supplement 1

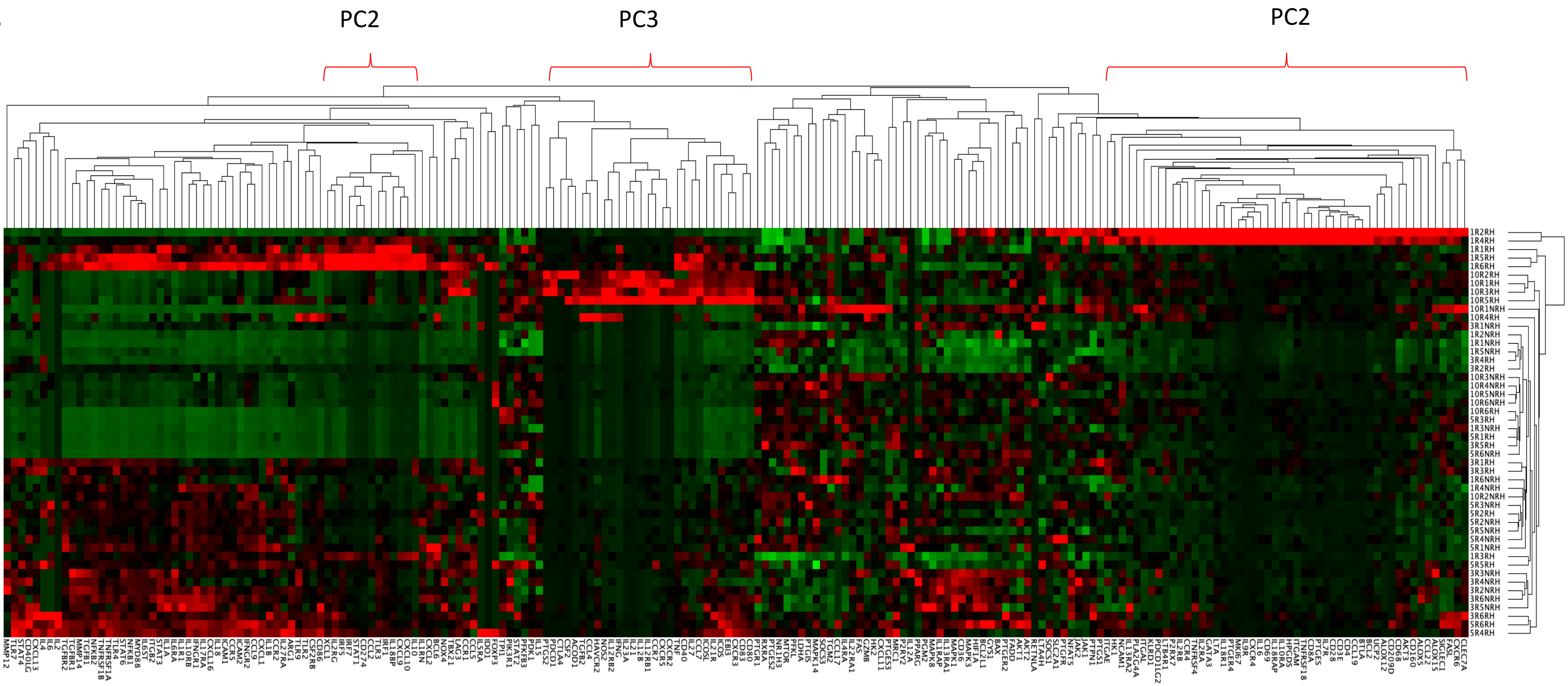

**Figure 2-figure supplement 1. Global statistical approach of gene expression - Unsupervised hierarchal clustering.** qPCR was carried out on mice liver tissue infected and control with L. infantum at 1, 3, 5 and 10 dpi. Unsupervised hierarchal clustering of the complete data: samples vs genes, was applied and based on Euclidean distance and pairwise average-complete linkage. For this representation was necessary autoscaling of NRQ data in *GenEx* software. Genes down-regulated during infection are shown in green and upregulated genes are in red.

**Figure 6- figure supplement 1**

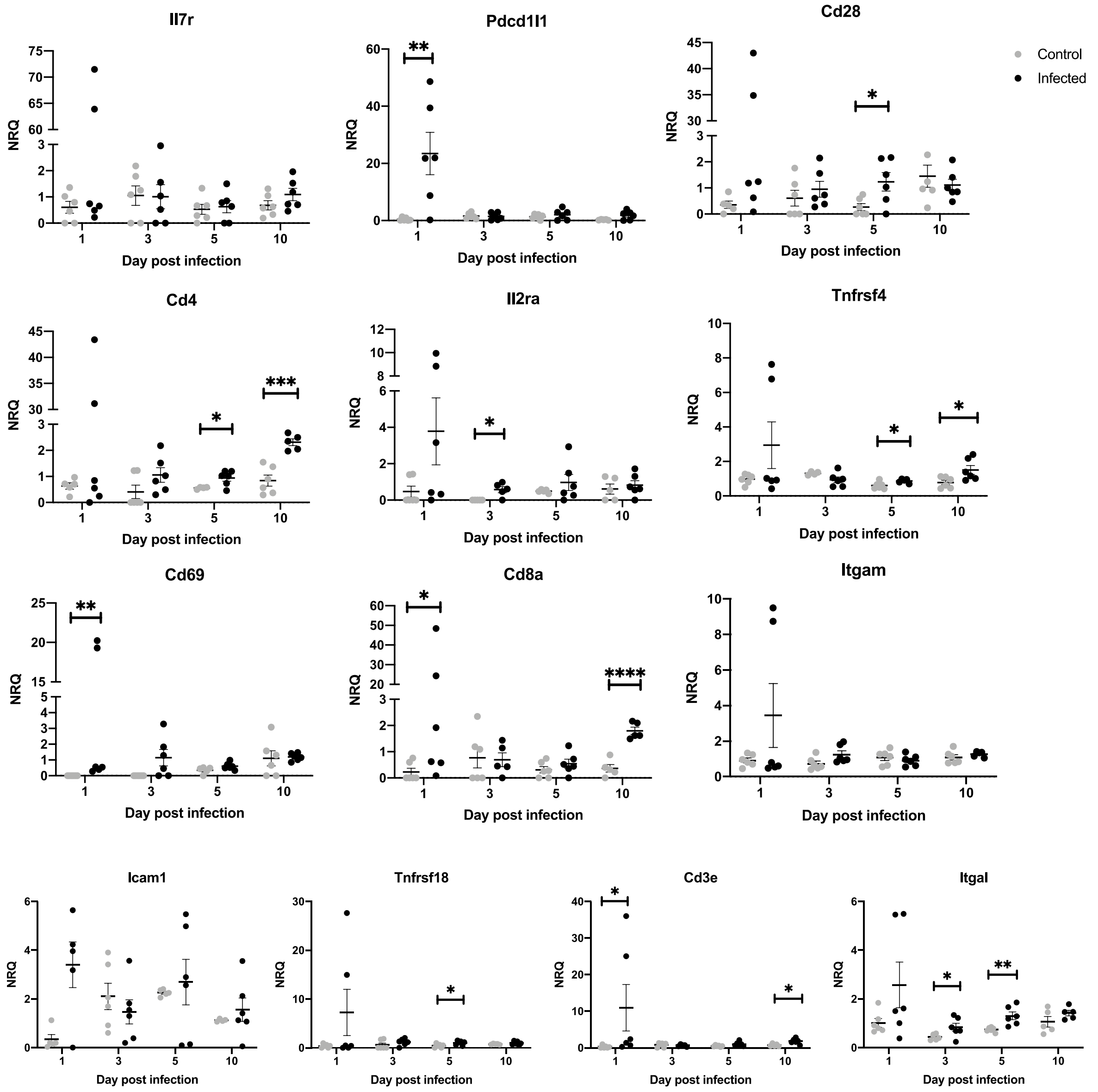

**Figure 6- figure supplement 1. Genes including in GO Component: External side of the plasma membrane.** Data are shown as the mean and SEM of the Normalized Relative Quantity values (NRQ). NRQ of Infected mice (black symbols) and control mice (gray symbols) are represented in scatter plot. Differences in NRQ values between non- infected (control) and infected animals were analyzed by two tail Mann-Whitney test or two tail unpaired t-test *p<0.05; **p<0.01; ***p<0.001.

### Figure 11- figure supplement 1

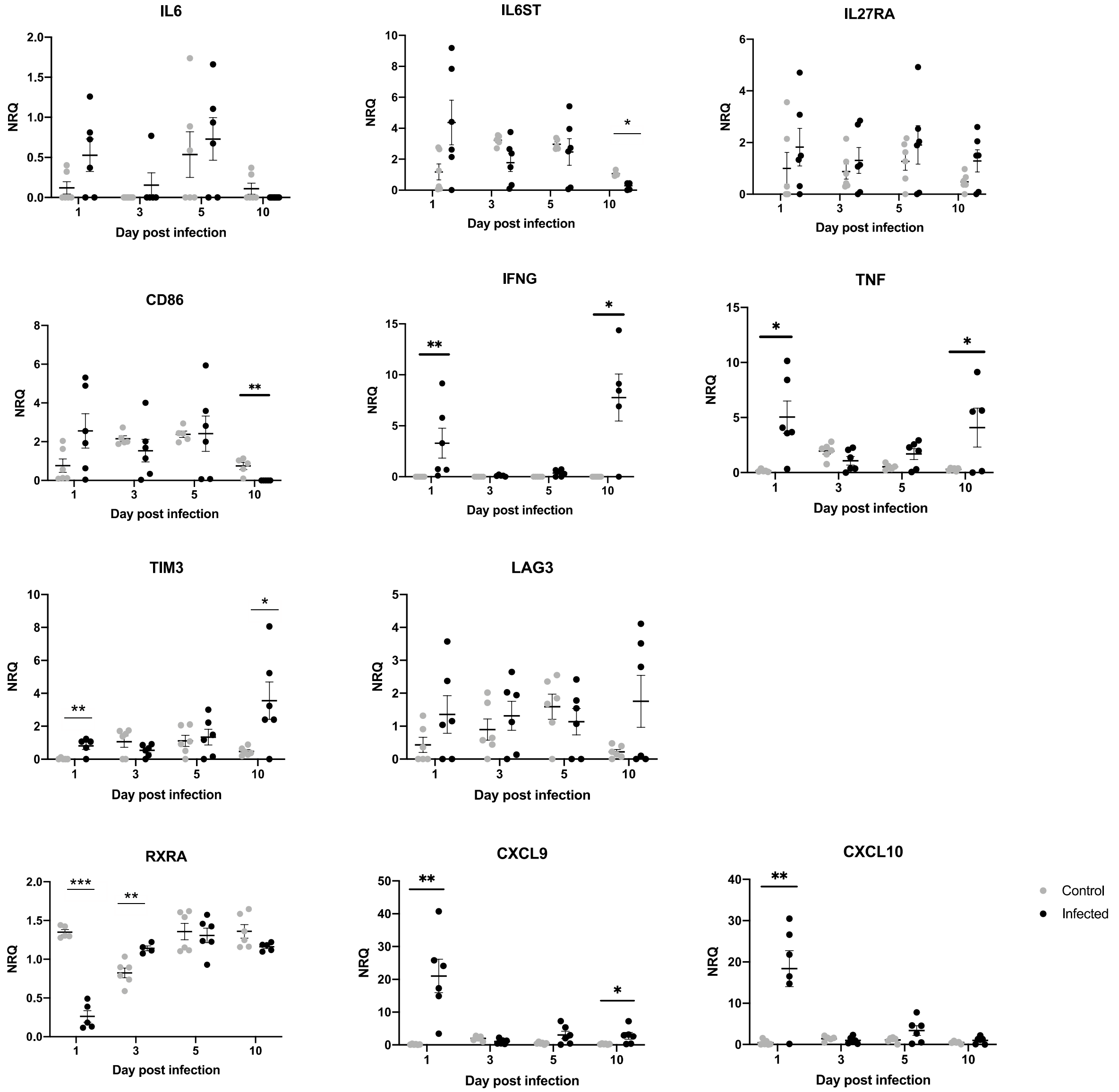

**Figure 11- figure supplement 1. mRNA expression of particular genes at 1, 3, 5- and 10-days post infection**. Data are shown as the mean and SEM of the Normalized Relative Quantity values (NRQ). NRQ of Infected mice (black symbols) and control mice (gray symbols) are represented in scatter plot. Differences in NRQ values between non- infected (control) and infected animals were analyzed by two tail Mann-Whitney test or two tail unpaired t-test *p<0.05; **p<0.01, ***p<0.01.
